# Breaking Down the Bottlebrush: Atomically-Detailed Structural Dynamics of Mucins

**DOI:** 10.1101/2024.04.09.588790

**Authors:** Fiona L. Kearns, Mia A. Rosenfeld, Rommie E. Amaro

## Abstract

Mucins, the biomolecular components of mucus, are glycoproteins that form a thick physical barrier at all tissue-air interfaces, forming a first line of defense against pathogens. Structural features of mucins and their interactions with other biomolecules remain largely unexplored due to challenges associated with their high-resolution characterization. Combining limited mass spectrometry glycomics and protein sequencing data, we present all-atom, explicitly solvated molecular dynamics simulations of a major respiratory mucin, MUC5B. We detail key forces and degrees of freedom imposed by the extensive O-glycosylation, which imbue the canonically observed bottlebrush-like structures to these otherwise intrinsically disordered protein backbones. We compare our simulation results to static structures observed in recent scanning tunneling microscopy experiments, as well as other published experimental efforts. Our work presents a generalizable framework for investigating the dynamics and interactions of mucins, which can inform on structural details currently inaccessible to experimental techniques.

## Introduction

Mucin (MUC) proteins comprise the primary macromolecular component of mucus, a viscous colloid secreted from, and with the fundamental role of protecting, the epithelium (**Figure 1A**). The MUC family of proteins is characterized by very long (100-1000nm, 200 kDa to 2.5 MDa) stretches called “mucin domains” that are rich in prolines, threonines, and serines, thus they are also referred to as “PTS” domains.^1–11^ PTS domains are composed of a linear protein backbone densely modified by O-glycans, resulting in over 80% of their total mass being comprised of carbohydrates (**Figure 1B-D**).^1–12^

**Figure 1:**
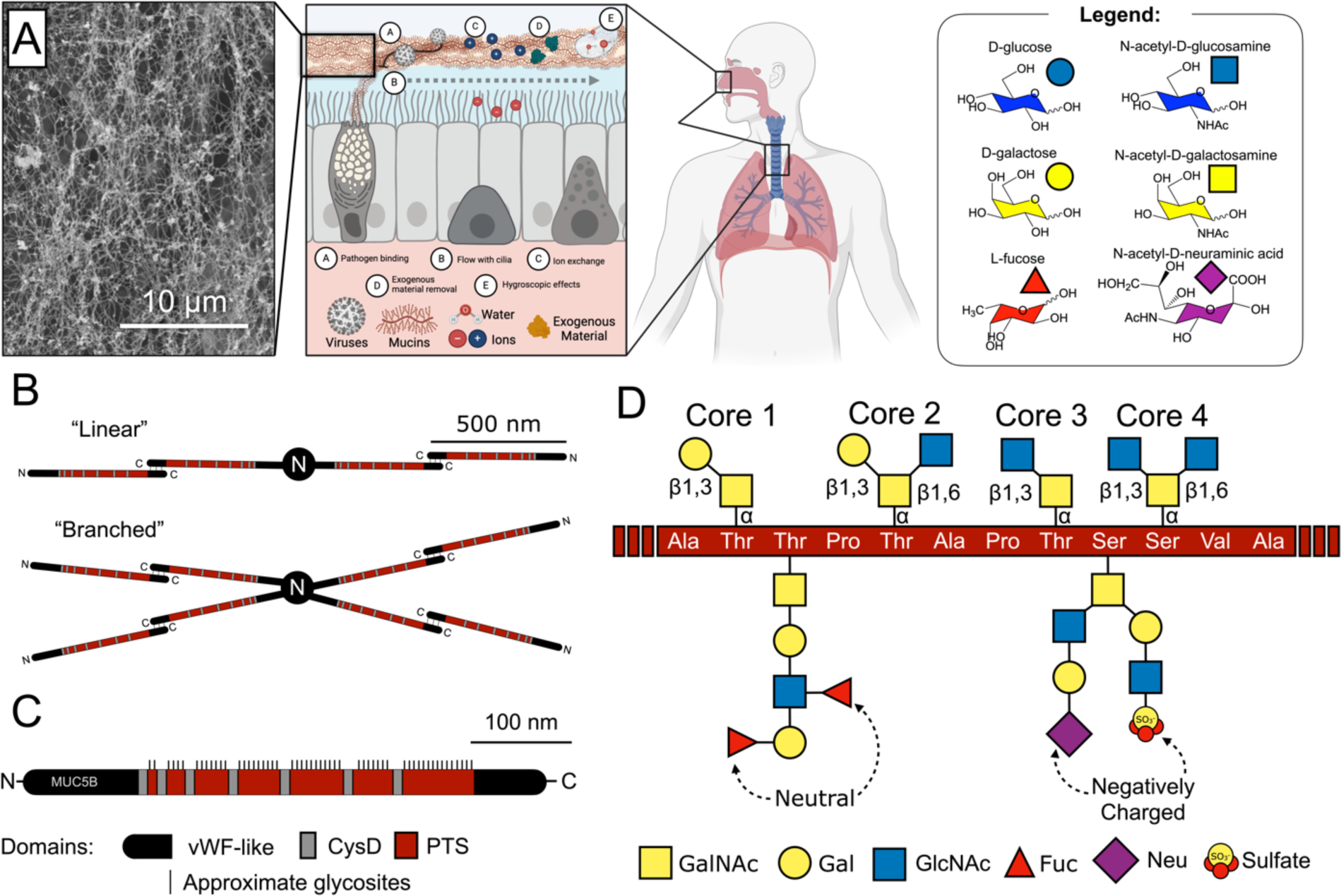
Legend detailing the saccharides discussed in this work and denoting their representations in symbol nomenclature for glycans (SNFG) (A) Schematic demonstrating the local of respiratory mucin expression, the many roles of mucins in the respiratory tract, and a scanning electron tomograph of a respiratory mucosal membrane, scanning image taken from Figure 1E in Carpenter et al.^86^ (B) Typical cross-linking schemes seen between mucin biopolymers, image adapted from Figure 1 in Symmes et al.^5^ (C) Structural domains of MUC5B mucin, image adapted from Figure 1 in Symmes et al.^5^ (D) Description of mucin structure at the atomic level, detailing common core types, connectivity, and ending motifs. All structures drawn with SNFG representation and color schemes; image adapted from Figure 1 in Symmes et al.^5^ Scheme in panel A created with Biorender.com.

Mucin-type O-glycans are characterized by modification of Ser and Thr residues (rarely also Tyr) with an initiating α-N-acetylgalactosamine (α-GalNAc). Following initiation, mucin O-glycans elaborate into several core structures and are often terminated by containing sulfate, sialyl, or fucose moieties (**Figure 1D**). While N-linked glycosylation has a characteristic amino acid motif (N-X-S/T) for glycosite identification, no such motif for O-glycans exists; instead, O-glycans can occur on any Ser or Thr residue within the PTS domain.^3,13–15^ Elaboration of O-glycan structure is non-templated and controlled by expression levels of glycosyltransferases, sulfotransferases, and sulfatases,^16,17^ all thus dependent on relative abundance of acceptor and donor substrates in the cellular milieu. Such factors make specific O-glycosite and O-glycotype prediction difficult to generalize.^3,18–25^ Overarching properties of mucin O-glycosylation, such as degree of charged sialyl and sulfate groups or neutral fucose groups, enable morphological responses to environmental changes such as pH and ion transfer.^4,26^ Furthermore, individual genetics and tissue type impact baseline expression of different mucins, and anthropological factors, such as smoking and disease status, impact O-glycan elaboration/branching/decoration.^5,9,27,28^ As a result, mucin domains demonstrate a great deal of macro-(is an O-glycan present at a glycosite or not), micro-(relative abundance of glycoforms found at each site including factors such as sialylation/sulfation/fucosylation), and meso-heterogeneity (as environmental factors such as pH and ion transport, or disease status can impart mucin morphological changes and/or up/down regulate certain O-glycoforms). Finally, protein backbones within mucin PTS domains are intrinsically disordered (**Figure 1B,C**),^29^ but such dense modification by O-glycans is canonically imbues a “bottlebrush”-like structure. Thus, O-glycosylation within the mucin domain impacts biophysical and biochemical forces on its surroundings modulated by ionic/pH microenvironment and expression levels.

The mucosal layer and its composite mucins play many roles in cell biology and physiology, including, but not limited to (**Figure 1A**): signal transduction, hygroscopic effects such as moisture maintenance, ionic exchange, and removal of pathogens or exogenous material through capture and subsequently moving in conjunction with ciliary beating.^1,5,8–13,16,17,30–44^ Such functions are dependent on the unique mucin structure, their resulting network-like mesh formed by long interacting strands (**Figure 1A**), and the ability to change morphology as a result of microenvironment ionic strength and pH.^4,26,43–45^ As mentioned, disease status can often impact mucin expression levels and O-glycoprofiles.^1,5,27,28,46^ For example, truncated aberrant mucin O-glycans are shown to be a hallmark of progression and metastasis in many cancers.^33,39–42,47^ Relatedly, one of the few FDA approved biomarkers for ovarian cancer is CA125, an epitope within the canonical mucin MUC16.^48–53^ Furthermore, increased expression of mucins are known to inhibit immune recognition either through physical barrier formation or inhibition of T-cells via binding of mucin sialic acids to Siglecs (sialic acid immunoglobulin-like lectins).^33,36,39–42,47^ Due to the many vital roles of mucins, dysregulation in expression, sequence, and structure can be detrimental to health and disease progression.

Despite their importance, because of their heterogeneity and their highly dynamic nature, structural characterization of mucins by traditional techniques such as cryo-electron microscopy and x-ray crystallography is often intractable.^29,34,54^. Mass spectrometry and glycoproteomics techniques, especially in combination with development of mucin active and specific protease (i.e., mucinases) enzymology, inform on the proportion of particular glycoforms and on which sites.^2,55–58^ Further, Nason et al. have elegantly demonstrated the ability to produce highly controlled mucin fragments^19,54^ which can then enabled Anggara et al. to, using scanning tunneling microscopy images, beautifully reveal images of MUC1 after gentle electrospray and surface deposition, representing some of the first work to “see” mucins in near single-molecule detail.^59^ However, there remains a gap between mucin glycoproteomic sequence information and high resolution structure and dynamics of mucins. Molecular modeling and simulation are beginning to fill these gaps. Recently, enabled by mass spectra acquired through characterization and upcycling of the mucinase SmE, we presented the first ever all-atom molecular dynamics (MD) simulations of a mucin-domain containing proteins, T-cell immunoglobulin mucin-domain containing protein 3 and 4 (TIM-3 and TIM-4).^60^ In that work, novel structural biology techniques in tandem with MD simulations revealed the roles that glycans play in the structure, stability, dynamics, and function of TIM-3 and TIM-4.^60^

In this current work, we describe the construction of mucin models for simulation using the respiratory mucin MUC5B as a test case. MUC5B is a massive (200 kDa to 2.5 MDa) glycoprotein that binds to and removes pathogens in the human airway, forming a thick physical barrier between our cells and the outside world.^4–6,27,29,31–34,37–39,47,61^ MUC5B is expressed and cleaved from goblet cells in the lungs, binds to pathogens and exogenous materials and removes them in conjunction with ciliary beating and aerosolization.^26,27,30,61–65^ MUC5B changes phase from gel-to fluid-like in the lungs as a function of ionic concentration (particularly Ca^2+^ and Cl^-^) and pH.^4,26^ We use published mass spectrometry glycoproteomic sequencing data to construct all-atom models of two repeating MUC5B respiratory mucin PTS domains (∼30 aa in length). We then scaled up this model by approximately an order of magnitude (∼224 aa in length) and conducted further simulation to explore structural dynamics at this second scale. Our simulations allow us to characterize atomic level forces and interactions driving the experimentally observed mucin “bottlebrush” morphology. Overall, in this work we: (1) describe a generalizable computational workflow for constructing mucin-domain containing glycoproteins, systems which are relevant to a vast array of biomedical and materials applications, (2) develop experimentally corroborated molecular models of MUC5B, and (3) predict atomic scale biophysical behavior driving macro- and mesoscale mucin morphology.

## Methods

With the goal to interrogate the atomic-level structural dynamics of respiratory mucins, we devised a workflow for constructing all-atom models of such species for simulation. We thus developed, constructed, and simulated two heterogeneous “mini mucin” models with slightly different degrees of charge. Our workflow centers on collecting relevant protein and glycomics sequence information (and/or choosing glycans from a database known to satisfy physiochemical properties, such as charge balance) and generating a small representative library for the work in question (**Figure 2**). Thus, while in many cases exact glycoforms cannot be resolved, representative and relevant molecular models of mucins can still be constructed to probe atomic scale properties.

**Figure 2:**
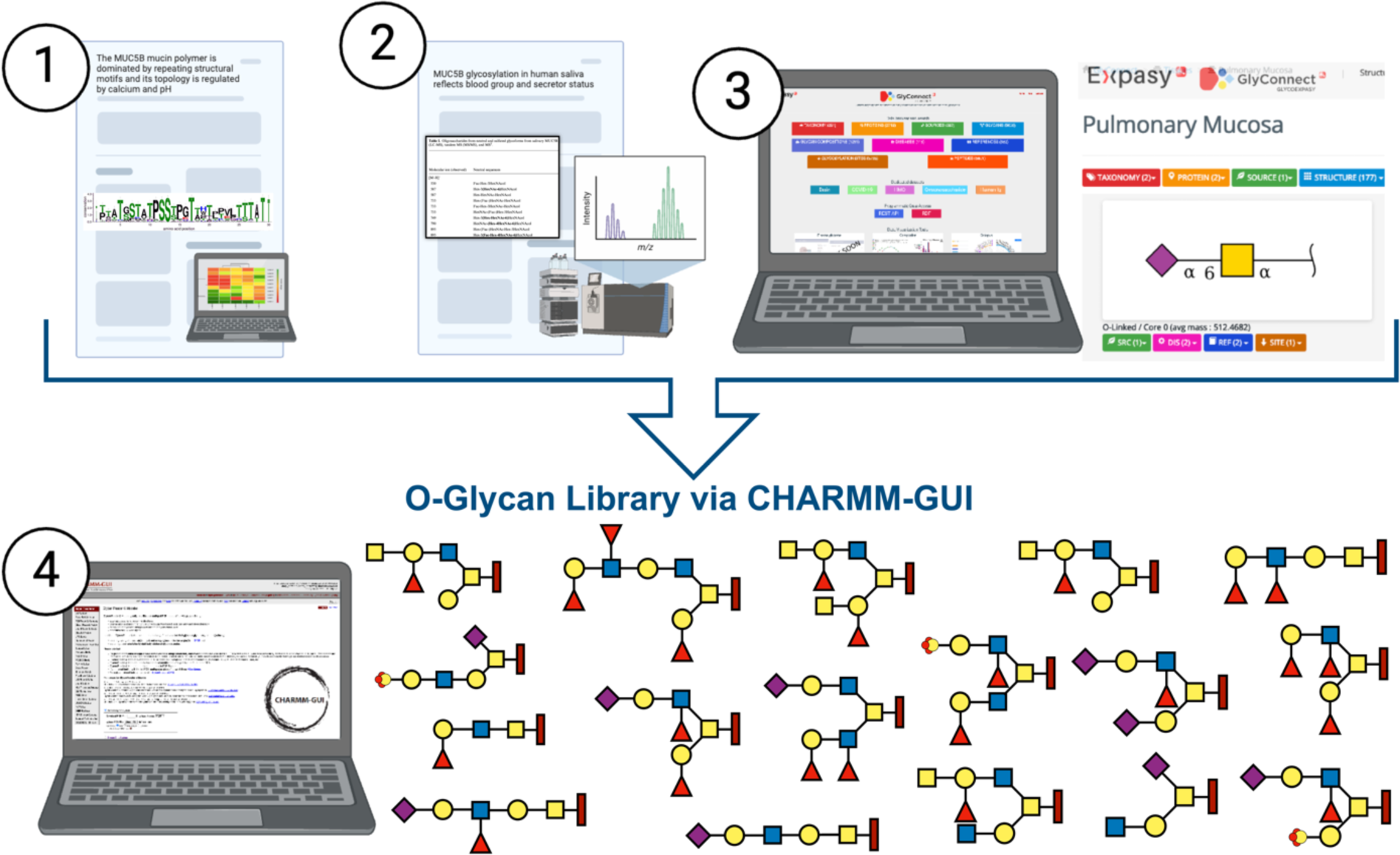
Workflow outlining how glycoproteomic data was collected to build miniature MUC5B mucin models simulated in this work. Steps 1 and 2 reflect collecting glycoproteomic data from literature resources. Step 3 reflects cross-checking those data with the Glyconnect Expasy Database and selecting O-glycan structures that are consistent with those data. Step 4 reflects constructing such O-glycans with CHARMM-GUI Glycan Builder. Created with BioRender.com.

We followed the workflow (**Figure 2**) to build two biochemically relevant models of short, ∼30 amino acid long, segments of MUC5B. Throughout this work, we refer to these two models as “Mini1” and “Mini2,” with each model reflecting a protein sequence based on a protein consensus sequence repeat within the MUC5B PTS domains. A complete step-by-step guide of our workflow can be seen in the supporting information (SI section 1.1.0) and we provide a summary of this workflow herein. Two protein MUC5B PTS domain consensus repeat sequences were taken from Hughes et al^5^ with amino acid (aa) lengths of 26 and 30, wherein they investigated the relative conservation of sequences within PTS domains.^4^ Rosetta’s BuildPeptide tool^66^ was used to build peptide structures of Mini1 (26 aa in length) and Mini2 (30 aa in length) sequences.^5^ We then compared extensive mass spectrometry (gel electrophoresis combine with LC-MS and LC-MS/MS) data from Thomsson et al.,^6^ which lists O-glycan structures from salivary mucosal samples, but without stereochemistry or connectivity information to O-glycans isolated in pulmonary mucosa from the GlyConnect Database with stereochemistry and connectivity information.^67^ O-glycans from the GlyConnect Expasy database were selected that corresponded to glycans with masses comparable to molecular ions listed in Thomsson et al’s^6^ Table 1. Additionally, care was taken to select a variety of charged and neutral glycans as well as higher proportion of Core-1 and Core-2 O-glycans than Core-3 and Core-4 O-glycans, as the former are slightly more common. CHARMM-GUI^68^ Glycan Modeller^69^ was used to add selected O-glycan structures from steps 2+3 to our Mini1 and Mini2 protein sequences. All Ser and Thr residues on Mini1+2 sequences were modelled with O-glycans (except Thr26 on Mini1, which was left unglycosylated). The positions of selected O-glycan structures were then “pseudo-randomized” onto their assigned mucin models: random numbers were used to generate initial positions, but positions were then modified to avoid over-clustering of charged glycans within mucins. The resulting O-glycoprofile maps for Mini1 and Mini2 can be seen in **Figure 3A** and proportions of core types and charged groups within Mini1 and Mini2 can be seen in **Table 1**.

**Figure 3:**
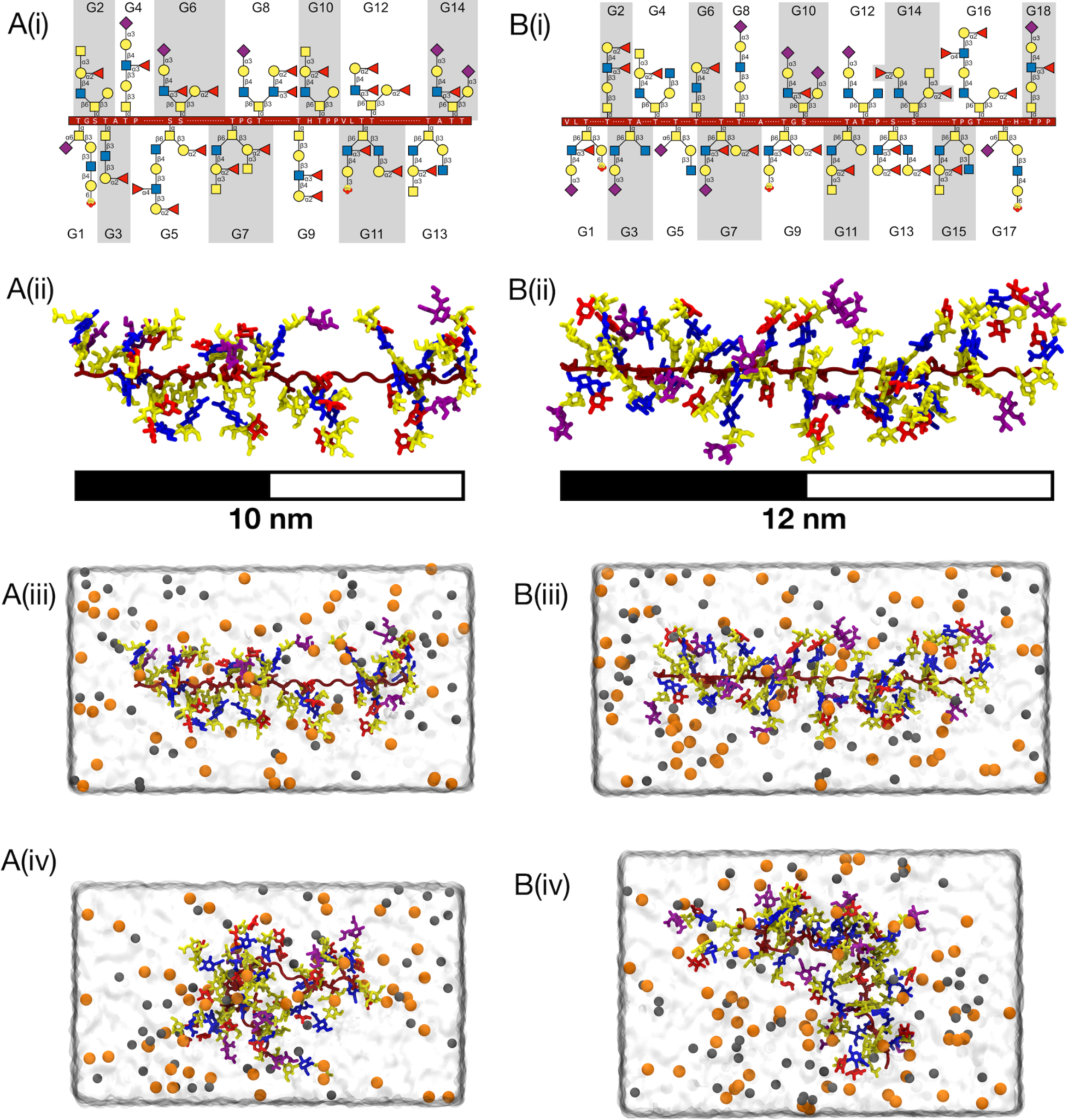
Construction of mini mucin systems. (A) Mini1 (i) glycoprofile and protein sequence, (ii) pre-minimization structure, (iii) solvated and neutralized structure, and (iv) snapshot from 550ns production simulations. (B) Mini2 (i) glycoprofile and protein sequence, (ii) pre-minimization structure, (iii) solvated and neutralized structure, and (iv) snapshot from 550ns production simulations. Protein backbones are colored in dark red, glycans are colored according to SNFG color scheme, Na+ and Cl-ions are shown in orange and grey spheres, respectively.

**Table 1:** Description of mass compositions in mucin models Mini1 and Mini2. Sulfate %mass calculated as mass of sum of SO^3-^ groups divided by total mass.

|  | Mini1 (#) | Mini1 (%mass) | Mini2 | Mini2 (%mass) |
| --- | --- | --- | --- | --- |
| #AminoAcids | 26 | 12.6 | 30 | 12.0 |
| #Glycans | 14 | 87.4 | 18 | 88.0 |
| #Core 1 | 3 | 16.8 | 5 | 21.8 |
| #Core 2 | 7 | 46.0 | 8 | 43.1 |
| #Core 3 | 2 | 10.3 | 1 | 3.0 |
| #Core 4 | 2 | 14.4 | 4 | 20.1 |
| #Sialylated Glyc (1,2) | 3, 1 | 8.9 | 8, 1 | 12.3 |
| #Sulfates | 2 | 0.8 | 3 | 1.0 |
| #Fucosylated Glyc (1,2,3) | 6, 5, 2 | 16.5 | 7, 3, 2 | 13.0 |
| Total Charge | -8 |  | -13 |  |

Additionally, to investigate the role of O-glycans in mucin conformational dynamics, i.e., in promoting a “bottlebrush” structure, we also constructed unglycosylated mini mucin systems using only the peptide backbones for each Mini1 and Mini2 **(Figure S1).** Following initial model construction of Mini1+2 glycosylated (**Figure 3B**) and unglycosylated **(Figure S1**) models, we then used VMD tools to solvate and neutralize these structures in TIP3 water boxes with 150 mM NaCl (**Table 2** lists system breakdowns for all mini models in both glycosylated and unglycosylated states). All simulations were performed in three replicas with NAMD3^70,71^ and parameters described by CHARMM36m^72–74^ all-atom force fields using San Diego Supercomputing (SDSC) resources. For Mini1 and Mini2 simulations: Periodic Boundary Conditions^75,76^ were applied, Particle Mesh Ewald^77^ was used to calculate long-range Coulombic potentials, Lennard-Jones potentials^78–80^ were calculated according to a 10 Å – 12 Å cutoff and switching scheme, pairwise list generation was cutoff at 15.5 Å. Langevin thermostat and piston were used to maintain temperature at 310 K unless otherwise stated and at 1 atmosphere of pressure. *<u>Minimization:</u>*To relax all conformations and alleviate close contacts resultant from CHARMM-GUI O-glycosylation steps, all solvated and neutralized structures were subjected to 10,000 steps of conjugate gradient energy minimization with no atomic restraints or constraints. *<u>Heating:</u>* Systems were then heated from 10 K to 310 K, increasing temperature by 25 K every 10,080 steps, timestep 2 fs, for a total of 240 ps of heating. Once at 310 K, an additional short equilibration of 770 ps of simulation was conducted. *<u>Equilibration:</u>*Mini1 and Mini2 were then subjected to NpT equilibration at 310 K for 0.5 ns, 2 fs timestep (useFlexibleCell option “on”, wrapWater option “on”, wrapAll option “on”). Systems were then subjected to an additional 55 ns NVT equilibration with 2 fs timestep with fixed box dimensions (useFlexibleCell option “off”). *<u>Production</u>*: Finally, 550 ns of NVT production runs were conducted at 310 K, timestep 2 fs. All data presented in the Results and Discussion section for glycosylated and unglycosylated mini mucin systems is derived from these 550 ns production runs.

**Table 2:** Description of total solvated and neutralized miniature mucin systems (glycosylated and unglycosylated) simulated in this work. Protein backbone lengths are listed for the fully outstretched state taken from the minimized frame.

|  | Mini1-Glyco | Mini2-Glyco | Mini1-Deglyco | Mini2-Deglyco | Mega-Glyco | Mega-Deglyco |
| --- | --- | --- | --- | --- | --- | --- |
| Total #atoms | 50,924 | 57,078 | 15,964 | 17,641 | 1,040,215 | 116,265 |
| #protein atoms/residues | 332 / 26 | 380 / 30 | 346 / 26 | 398 / 30 | 2827 / 224 | 2899 / 224 |
| Protein Backbone Length (Å) | 88 | 99 | 88 | 99 | 769 | 769 |
| #glycan atoms/#glycans | 1,055 / 14 | 1,143 / 18 | --/-- | --/-- | 20,492 / 128 | --/-- |
| #water atoms/residues | 48,231 / 16,077 | 53,841 / 17,947 | 15,588 / 5,196 | 17,211 / 5,737 | 506,343 / 168,781 | 113,154 / 37,718 |
| #Na/#Cl ions | 53 / 45 | 64 / 51 | 15 / 15 | 16 / 16 | 518 / 450 | 106 / 106 |
| System dimensions (ÅxÅxÅ) | 63.9 x 68.7 x 124.99 | 65.7 x 68.4 x 137.1 | 36.2 x 39.6 x 120.8 | 36.7 x 40.0 x 134.6 | “ “ | “ “ |
| Total Sampling | 3 x 550 ns | 3 x 550 ns | 3 x 550 ns | 3 x 550 ns | 210 ns, 283 ns, 264 ns (forked) | 3 x 150 ns |

To begin to approach more physiologically relevant mucin conformations, we also simulated larger “Mega” mucin models. We constructed our “Mega” mucin system by patching the C-terminal of Mini1 to the N-terminal of Mini2, then quadrupling this Mini1-Mini2 unit to form a long chain (all done with VMDtools,^81^ see shared files and Supporting Information). The final resulting Mega-mucin was 769 Å in length along the protein backbone, comprising 224 amino acids with 128 glycans. As with the Mini mucins, we also sought to investigate the impact of O-glycans on mucin backbone conformations, thus we constructed a unglycosylated Mega mucin model by removing O-glycans and generating resultant psf/pdb starting files with psfgen. Similar to the mini mucin systems, the mega mucin glycosylated and unglycosylated systems were solvated and neutralized in a TIP3 water box with 150 mM NaCl, minimized, heated, and equilibrated according to a similar standard simulation protocol. Due to the sheer size of the Mega mucin system (>1 Million atoms, **Table 2**), we forked the simulation to optimize computational usage and acquire robust sampling. To do this, we performed an extended equilibration of 150ns and extracted a frame to restart three replicates at reinitialized velocities. The glycosylated Mega mucin system was then production simulated with NAMD3^70,71^ using SDSC resources for an aggregate of 757 ns, while the unglycosylated Mega mucin was production simulated with NAMD3 for an aggregate of 450 ns.

Following the above construction and simulation protocols, we then conducted extensive analysis of these simulations using MDAnalysis and in-house scripts,^82,83^ see Supporting Information files for all analysis scripts. In several of the analyses we refer to relative values, such as relative end-to-end distances. These relative values are calculated as ratios of, e.g., the end-to-end distance per frame by the end-to-end distance of the minimized (outstretched) structure before simulation. Thus, in this example, a relative end-to-end distance value approaching 1 would indicate the protein backbone in that frame is nearly as extended as the minimized state, whereas a value around 0.5 indicates the two ends of the protein backbone is half as extended as the minimized state, and less than 0.5 indicates the two ends of the protein backbone are very close to each other, as would be required for a compact conformation.

## Results and Discussion

### Characterizing biophysical properties for glycosylated mini mucin models

We sought first to compare glycosylated Mini1 versus Mini2 simulations to identify if the constructed O-glycan and protein sequence heterogeneity induced significant dynamical differences. No significant differences between Mini1 and Mini2 were exhibited in the following properties: persistence length, relative end-to-end distances, relative radius of gyration (Rg, ratio of instantaneous Rg to minimized Rg), relative compactness (ratio of instantaneous compactness relative to minimized compactness), curvature per C_α_, angle between each C_α_, and Ramachandran and Janin plots for glycosites and non-glycosites. All Mini1 vs. Mini2 glycosylated data can be seen in the Supporting Information **Figure S4**, for complete details on how these data were calculated please see SI Methods Section 1.3. These calculations indicate that, despite their specific protein and O-glycan sequence differences, on a global level the mini mucins demonstrate markedly similar conformational landscapes. Significant differences were observed for root mean square fluctuations (RMSFs) on a per-residue basis between Mini1 and Mini2 and we discuss those differences in detail later.

To better characterize the conformational landscapes accessible to Mini1 and Mini2 glycosylated mucin structures, we conducted principal component analysis (PCA) on all backbone conformations from all Mini1 and Mini2 simulations (3 replicas of 550 ns each, all trajectories concatenated and aligned by C_α_s, Supporting Information **Figure S2**). PCA revealed 3 PCs could explain greater than 60% of variance in C_α_ atomic positions. We clustered all frames according to these 3 PCs with scikit-learns kmeans clustering module to identify 11 clusters. Clusters were identified via sum of square error inflection point and silhouette coefficient. See SI Methods Section 1.3.1 for complete details about how trajectories were merged and how PCA and kmeans clustering were conducted. Per cluster, we then calculated a relative linearity score (RLS) according to Equation 1, below:

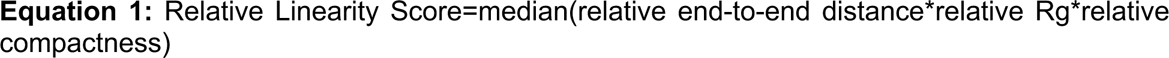

Details on how each of these terms were calculated per frame is shown in the Supporting Information Methods Section 1.3. We then ordered the original clusters from 0-10 according to this RLS for ease of comparison: 0 having the lowest RLS (which estimates a “low linearity”, thus “high curvature”/“high compactness”), 10 having the highest RLS (which estimates a “high linearity“/”low compactness”). Protein backbone conformations for clusters 0-10 can be seen in **Figure 4A**, alongside relative populations for each cluster in percentage. Each cluster’s bar is colored according to that cluster’s RLS. Distributions of all terms required for RLS calculation per cluster are shown in Supporting Information **Figure S3.** We did see that some clusters were more populated for glycosylated Mini1 or Mini2 systems. For example, clusters 3, 4, and 8 were highly popular for Mini1, while clusters 2, 5, 6, and 7, were highly popular for Mini2. However, representative structures for these “medium” linearity clusters are similar and distinctions between cluster populations are likely due to sampling limitations. Thus, at this time, we cannot identify glycoprotein sequence differences that would lead to conformational landscape preferences. We did see that the cluster with the highest curvature, Cluster 0, was predominantly populated by Mini2, but, while not highly populated, Mini1 did also produce some frames in this cluster.

**Figure 4:**
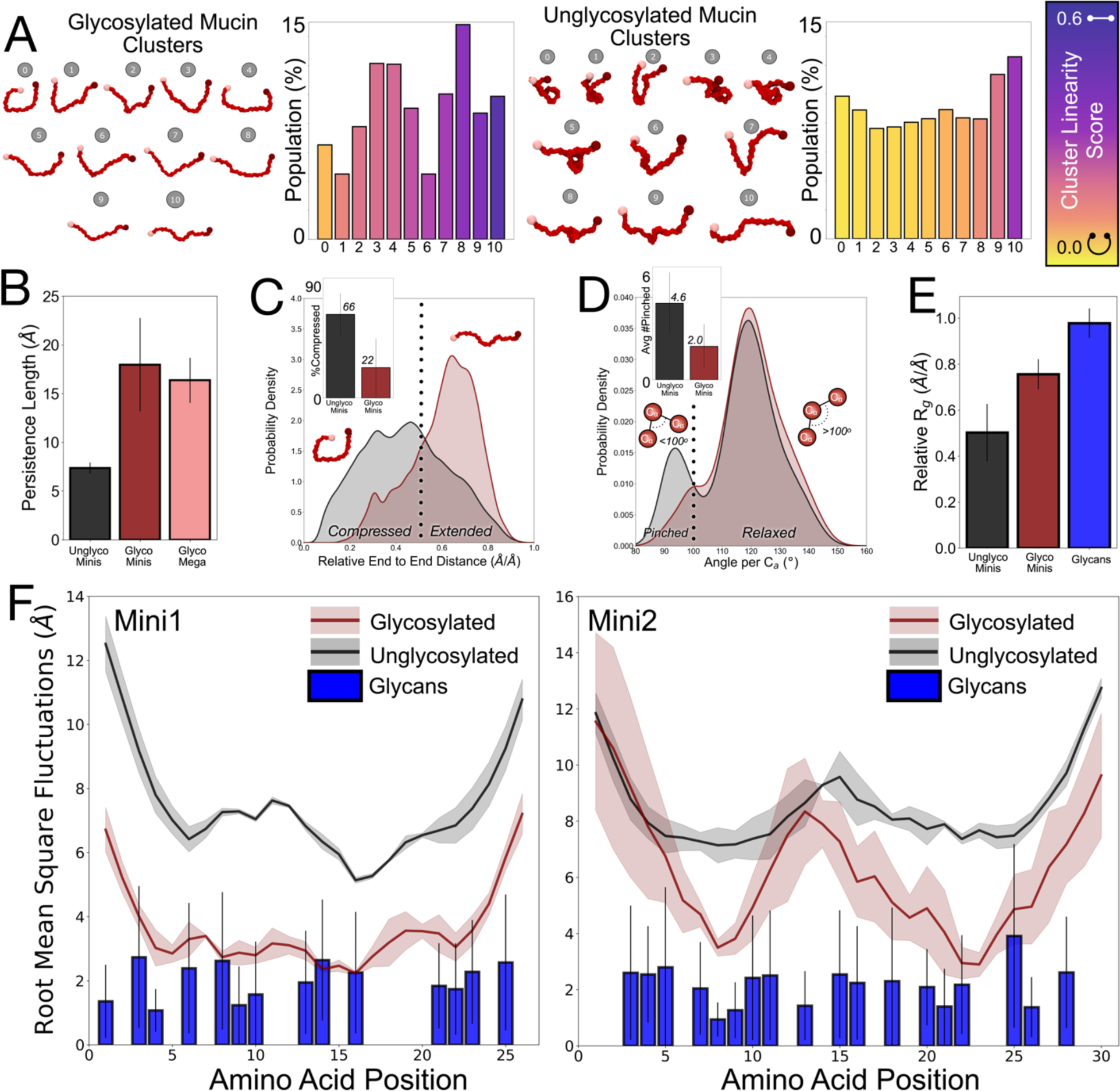
Describing the bottlebrush. (A) Representative structures from glycosylated and naked mini mucin clustering results alongside cluster population densities (%). Cluster population bars are colored on a scale according to cluster linearity score. In (B-F) black, red, pink, and blue trends/bars indicate naked mini mucin simulations, glycosylated mini mucin simulations, glycosylated mega mucin simulations, and glycans from glycosylated mini mucins simulations respectively. (B) Persistence lengths calculated from naked mini mucins, glycosylated mini mucins, and glycosylated mega mucins. (C) Relative end to end distances calculated for glycosylated mini mucins and naked mini mucins. Inset depicts the percent of frames naked and glycosylated mini mucins spend in “folded” conformations, relative end to end distance less than 0.5. (D) Distributions of angles per C_α_ for glycosylated and naked mini mucins. Angles per C_α_ less than 100° correspond to rotations consistent with alpha helices and are thus marked “pinched”. Inset depicts the average number of “pinched” C_α_ per frame in naked and glycosylated mini mucins. (E) Average relative radius of gyration calculated for naked mini mucins, glycosylated mini mucins, and for all glycans from glycosylated mini mucin simulations. (F) RMSFs calculated for naked and glycosylated Mini1 and Mini2.

Mini1 and Mini2 were constructed with different protein backbones and O-glycoprofiles, and yet they have remarkably similar conformational landscapes and biophysical properties, as seen in the clustering results and persistence lengths (Supporting Information **Figure S2-4).** Although the two mini mucin sequences are distinct, they were both constructed from MUC5B protein consensus repeat sequences and pulmonary mucosa O-glycoforms found within the Glyconnect Expasy database. That the two resultant models behave similarly suggests that key biophysical characteristics of mucins are stored within these consensus sequence repeats. Furthermore, we hypothesize from these results that although there is a great deal of O-glycan heterogeneity within mucin domains, if the overarching parameters are maintained – i.e., type similarity in protein backbones, O-glycan density, and charge density – then backbone conformations, and overall mucin morphology are likely similar.

### Mucin models adopt distinct conformational states in glycosylated and unglycosylated states

To probe how mucin O-glycans promote a bottlebrush-like structure on mucin protein backbone conformations, we compared all glycosylated mini mucin data to all unglycosylated mini mucin data. Several groups have characterized the bottlebrush through electron microscopy and other techniques and have begun to piece together the key forces related to the nature of the “bottlebrush”.^4^ Our MD simulations reveal that the mucin O-glycan bottlebrush results in increased linearity of backbone conformations, increased persistence length along the peptide backbone, and decreased flexibility of backbone residues.

In addition to the clustering performed for the glycosylated mini mucins described above, we also performed PCA guided kmeans clustering for all unglycosylated mini mucin simulations (Supporting Information **Figure S5**). Here we see a significantly different array of representative cluster shapes. Comparing clusters between unglycosylated and glycosylated mini mucin simulations, unglycosylated clusters are far more compact, non-linear, and compressed with scores ranging from 0.03 to 0.41 with a median score of 0.15. In contrast, glycosylated mini mucin clusters have linearity scores ranging from 0.08 to 0.51 with a median of 0.48. Indeed, for unglycosylated mini mucin simulations, 9 out of 11 clusters have low linearity scores (<0.2) indicating compressed states, while only 5 out of 11 glycosylated mini mucin clusters have such low linearity scores. Despite their propensity to adopt more compressed or curved clusters, unglycosylated mini mucin simulations did still populate linear conformations with unglycosylated cluster 10 having a comparable linearity score to the glycosylated cluster 10, 0.41 and 0.52 respectively (Supporting Information **Figure S6**). In addition, while compressed conformations were far more populated for unglycosylated mini mucins (76.1% of frames in clusters with linearity scores <0.2), unglycosylated mini mucins can still adopt conformations like the linear conformations seen in the glycosylated mini mucin simulations (24.0% of unglycosylated frames in clusters with >0.3 linearity scores, 12.6% of frames in cluster 10 with 0.41 linearity score). Taken together, these results indicate that O-glycans impose steric limitations that shift the mucin conformational landscape to more linear backbone structures. Our data suggest that O-glycans prevent compression of the mucin protein backbone and thus prevent accessing more compact shapes.

For each simulation – glycosylated, unglycosylated, mini, and mega -- we also calculated the persistence length (PL) along the mucin protein backbone and observed a drastic difference in PLs between unglycosylated and glycosylated models: unglycosylated mini simulations with a PL of 7.3± 0.6, glycosylated mini simulations with a PL of 17.9 ± 4.8, (**Figure 4B** and Supporting Information **Figure S7).** As described, there was marked agreement between PLs calculated for glycosylated mini systems and glycosylated mega systems, 17.9 ± 4.8 versus 16.4 ± 2.3, respectively. Thus, the presence of O-glycans along mucin backbones increases PL by a factor of 2-3 compared to unglycosylated systems. We also tracked relative end-to-end distances, which reveal a dramatic difference in the distribution of ratios between unglycosylated and glycosylated mini mucin systems. Unglycosylated mini mucins demonstrate a broad distribution of states from fully extended (relative end-to-end distance ∼0.8) to fully compressed (relative end-to-end distance ∼0.2) with a mean of 0.44±0.19. In contrast, glycosylated mini mucins demonstrate a sharper distribution of states with a mean of 0.60±0.15. Unglycosylated mucins spend ∼62.3%±15.9% of frames in compressed conformations whereas glycosylated mucins spend only ∼21.4%±18.0% of frames in such (**Figure 4C**, where we define compressed conformations defined as having a relative end-to-end distance <0.5).

Upon observing representative structures from clustering, as well as observing trajectory frames, we noted many simulation conformations adopted “curved” and “pinched” structures, with significant backbone orientation changes around specific protein residues. For example, see the representative structure glycosylated mini mucin cluster 2: in this representative structure there is at least one distinct “pinch point”. To quantify the probability of each amino acid along each mini mucin (glycosylated and unglycosylated) adopting such a “pinched” conformation, we calculated the distributions of angles formed between each C_α-1_, C_α_, and C_α+1_ (excluding terminal C_α_s), **Figure 4D**. We see that unglycosylated mini mucin systems have a distinct global and distinct local energetic minimum (population maxima) for C_α_ angles, while glycosylated mini mucin systems have a distinct global energetic minimum, with a smaller shoulder in overlapping with the local minimum seen in unglycosylated simulations. α-helical turns are characterized by ∼100° rotation per amino acid, i.e., each complete turn of the alpha-helix is occupied by 3.6 amino acids. This ∼100° rotation angle corresponds to the local energetic maximum wherein unglycosylated mini mucin C_α_s have decreased population. As such we demarcated ∼100° as a “pinch point”: C_α_s with conformations below 100° were to be considered “pinched.” We then calculated the average number of pinched C_α_s per frame: unglycosylated mini mucins exhibit an average of 4.6 ± 1.8 pinched C_α_s while glycosylated mini mucins exhibit an average of 2.0 ± 1.3. Thus, unglycosylated mini mucins demonstrate nearly 2x more pinched C_α_s than glycosylated mini mucins and, on average, have enough “pinch points” to generate nearly circular backbone conformations. Per residue distributions of C_α_ angles identify residues which are more/less likely to adopt a pinched conformation. Such pinching conformations in glycosylated mini mucin simulations are more likely for non-glycosite residues and residues outside of densely glycosylated regions (Supporting Information **Figures S8,S9),** as these residues on average have broader C_α_ angle distributions or bimodal distributions suggesting distinct conformational states.

To isolate which components within each mini mucin structure are relatively more or less rigid, we calculated root mean square fluctuations (RMSFs) and relative Rgs for protein residues and glycan segments separately. To calculate relative Rgs, we calculated first the radius of gyration for the minimized (outstretched) protein back bone, and then for each frame of the simulation we tracked the ratio of the framewise Rg to the minimized Rg. To calculate the glycan relative Rg, we similarly calculated the Rg for each glycan in the minimized structure, and then per-frame calculated the ratio of framewise Rg to the minimized radius of gyration. From these relative Rgs, we see that each glycan is relatively rigid with a sharp distribution in relative Rgs around 0.98± 0.06, **Figure 4E**, see also supporting information **Figure S10** for per glycan average relative Rg. The distribution of relative Rgs for the glycosylated mucins was a bit broader, but still high, with an average of 0.76± 0.07. However, the distribution of relative Rgs for unglycosylated mucins is much lower and broader, with an average of 0.50±0.13. Furthermore, from RMSF calculations, we see that glycosylated mucin protein backbones as well as O-glycans are far more rigid relative to their unglycosylated counterparts, **Figure 4F**. Additionally, some O-glycans are less flexible than the protein backbone in glycosylated mucin simulations. All our results underscoring the rigidity and linearity of mucin protein backbones are supported rigorously by Kramer et al^84^ who demonstrated the increased rigidity of mucins as a function of O-glycan density. Our results paint hitherto unseen details of the mucin “bottlebrush” structure: dynamics of a flexible, otherwise intrinsically disordered protein backbone, are dampened by the rigid/inflexible O-glycans.

### Detailing the physical characteristics of a large-scale bottlebrush

To understand the relationship between mucin model length and the nature of the “bottlebrush”, we constructed a larger, more biologically relevant Mega mucin model. The resulting structure was 224 amino acids in length with 128 O-glycans. While this model is still far from the true scale of MUC5B, which can be as large as 6000 amino acids long, simulations of this longer model allow us to explore conformational states accessible to longer length mucins. As done with the Mini mucins, we also constructed a unglycosylated Mega model with the same protein sequence but no O-glycans. The glycosylated Mega mucin adapted an almost equal distribution of compressed and extended conformations (**Figure 5A**), whereas the protein backbone in the unglycosylated Mega systems took on largely compressed conformations (<0.5 relative end-to-end distance, ∼91% of population) with normalized end-to-end distances concentrated close to a ∼0 relative end-to-end distance. The conformational differences between glycosylated and unglycosylated Mega mucin systems are expanded upon in the conformational clustering done on the resultant trajectories (see Supporting Information **Figures S11, S12** for details on how clustering was performed for all mega mucin simulations). The unglycosylated and glycosylated Mega mucin clusters (**Figure 5B** and **5C**, respectively) are numbered from most to least occupied, with 1 being the highest populated cluster and 5 being the least populated cluster. Consistent with the normalized distance plot, the most occupied cluster for the unglycosylated Mega mucin (54%) is that of a fully compact protein backbone, completely collapsed on itself. By comparison, glycosylated Mega mucin clusters are all relatively extended, **Figure 5C**. Thus, these results suggest the O-glycans impose structural or conformational rigidity along the protein backbone.

**Figure 5.**
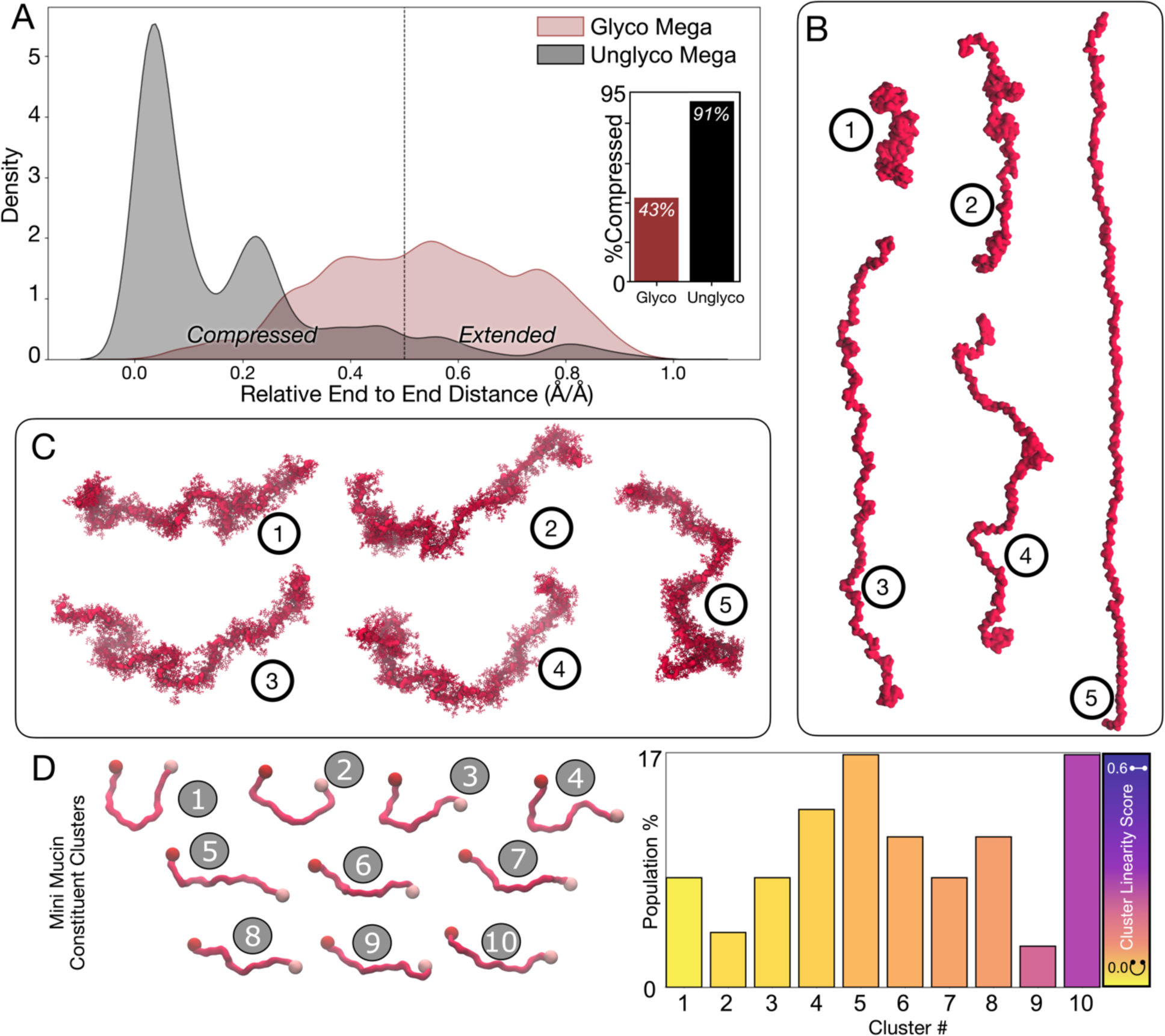
Relationship between size, glycosylation, and compactness of mega mucins. (A) Relative end-to-end distances calculated for glycosylated mega mucin and unglycosylated mega mucin. Inset details percent of the respective populations observed in “folded” state (i.e. > 0.5 normalized distance) vs. “unfolded” state (i.e. <0.5 normalized distance). (B) Representative structures extracted from cluster centroids resultant of unglycosylated mega mucin simulations. Cluster numbers 1, 2, 3, 4, and 5 represent highest to lowest observed population – 54%, 23%, 10%, 9%, and 4%, respectively. (C) Representative structures extracted from cluster centroids resultant of glycosylated Mega mucin simulation. Cluster numbers 1, 2, 3, 4, and 5 represent highest to lowest observed population – 28%, 24%, 19%, 16%, and 13%, respectively. (D) Representative structures from mini constituent mucin clustering alongside population densities (%). Cluster population bars are colored on a scale according to cluster linearity score.

Our Mega mucin clustering results also highlight the unique balance between order and disorder within the mucin domain. As seen in the representative structures from glycosylated Mega mucin clustering (**Figure 5C**), each cluster centroid looks qualitatively similar, and cluster population distribution is evenly distributed (28%, 24%, 19%, 16%, and 13%), however there are subtle enough differences to be clustered differently in PCA space by kmeans clustering (Supporting Information **Figure S13**). What’s more, although the global shape of each structure looks similar, differences can be spotted between structures related to where bends occur along the protein backbone. We can account for the overall qualitative similarities despite their minute differences by recalling that mucins are comprised of an intrinsically disordered protein backbone core which is densely modified by O-glycans. The protein core provides underlying dynamic flexibility, but that flexibility is modulated by steric-bulk imparted by the rigid O-glycans (Supporting Information **Figure S14**). Thus, while glycosylated mucins cannot adopt fully compressed conformations as seen in the unglycosylated simulations, they can adopt a multitude of similarly curved conformations. Further investigation will probe the degree to which O-glycan sequences alter the positions of these backbone curves and the functional impacts such curves may serve, such as pathogen binding.

To identify if the ∼30 amino acid peptides within our Mega mucin model were also adopting conformations similar to those exhibited by the Mini mucin models, we characterized several parameters from each of the clusters and screened each 26aa window along the mega mucin backbone to identify which clusters from Mini mucin simulation were most like that of conformations generated from the Mega mucin (**Figure 5D**). Interestingly, the most observed constituent Mini mucin conformations derived from the mega mucin simulation accessed fewer distinct conformations than Mini mucins simulated in bulk and had markedly consistent linearity scores across all clusters. This result suggests that constituent Mini mucin segments within the Mega mucin model are more conformationally constrained by the steric effects of neighboring mini mucin constituents to which they are bonded. Although there is conformational overlap between the Mini mucin constituents of the Mega mucin and the bulk Mini mucins, this analysis demonstrates that mini mucins in bulk may over-sample a conformational landscape inaccessible to larger, more physiologically relevant, mucin models. However, we did observe significant overlap for the constituent Mini mucins and the bulk Mini mucins according to several physical characteristics: the average linearity score for the constituent mini mucins is similar to the average linearity score for the mini mucins simulated in bulk. Additionally, we calculated nearly the same persistence length for glycosylated Mini mucins and glycosylated Mega mucin models (**Figure 4B**). Thus, for future simulations wherein one aims for physiologically consistent mechanical and chemical properties, such as PL, and not looking to extract exact conformational landscapes of global mucins, mini mucins simulated in bulk are likely sufficient.

### Elucidating the forces that imbue structure within the bottlebrush

We next sought to characterize the biophysical relationships between protein and O-glycan residues that result in pseudo-structural properties within the bottlebrush domain. To do so we tracked several degrees of freedom from Mini glycosylated and unglycosylated simulations. Firstly, we calculated Ramachandaran and Janin plots for mucin models in glycosylated and unglycosylated states, and we plotted those degrees of freedom for non-glycosites (i.e., residues that are not glycosylated in glycosylated simulations, thus not Ser and Thr residues) and glycosites (i.e., Ser and Thr residues that *would* be glycosylated in glycosylated simulations, but which are not glycosylated in unglycosylated mucin simulations), **Figure 6A**. These results demonstrate a stark difference in torsional degrees of freedom for the protein backbone in glycosylated and unglycosylated states. We see that glycosite residues in glycosylated mucin simulations have a loss of probability density for Ψ backbone torsions in the range of ∼-120° to ∼-50° compared to those same residues in the unglycosylated simulations and compared to non-glycosites in glycosylated simulations. We see that from the glycosylated simulations, non-glycosites occupy Ψ values in the range of −180° to - 60° in ∼10% of frames. However, in these same simulations, glycosites do not occupy this state (calculated through area under the curve across Ψ kernel density function, see supporting information **Figure S15** for 1D densities). Similarly, non-glycosites occupy the −60 to 60 Ψ domain in approximately 16% of frames, while glycosites occupy this domain in only ∼6% of frames. Finally, glycosites largely populate the 60 to 180° range of Ψ angles, at 92% population versus 72% for non-glycosites. Thus, O-glycosylation dramatically reduces the degrees of freedom accessible to mucin protein backbone conformations.

**Figure 6:**
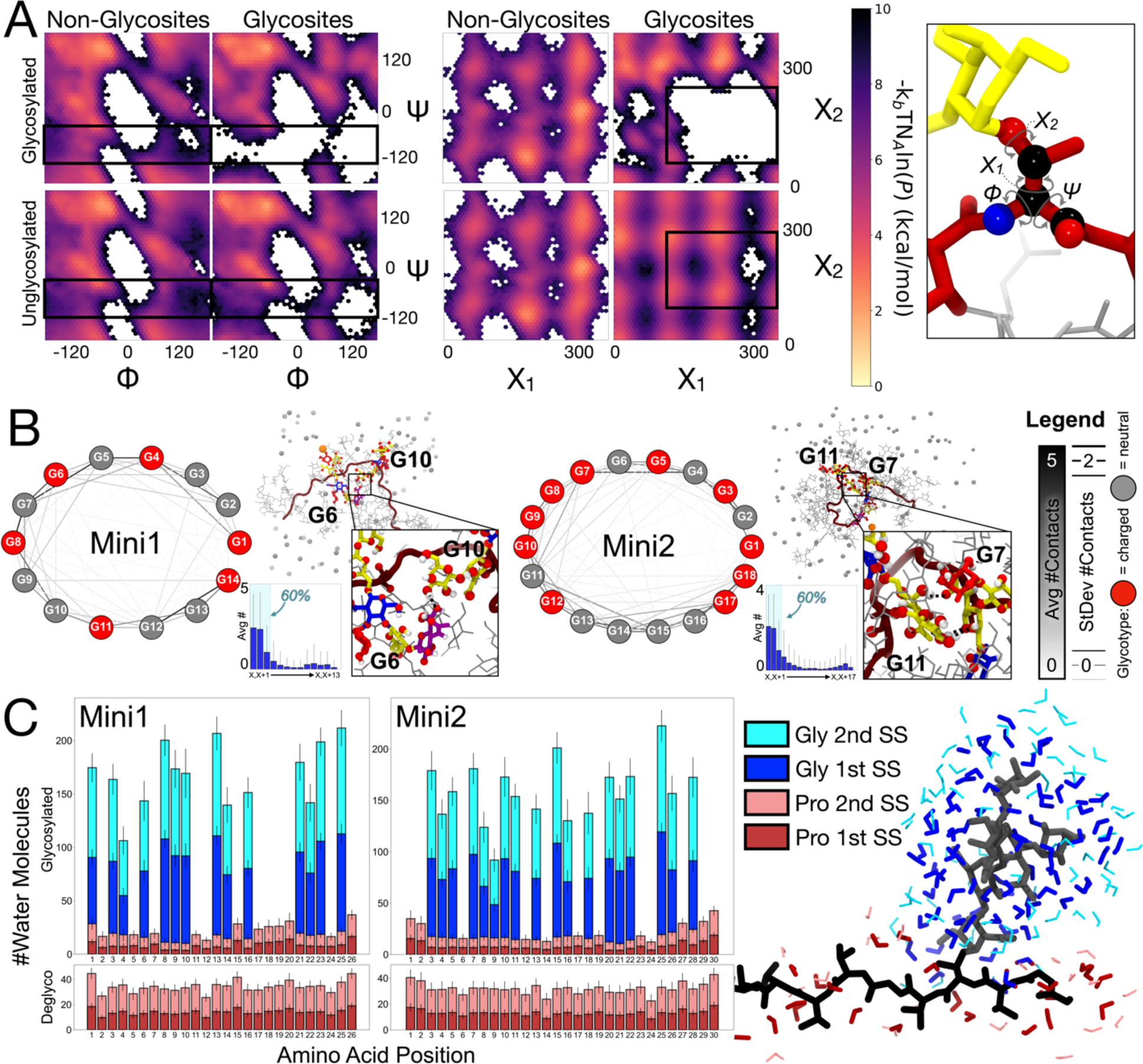
Elucidating the bottlebrush. (A) Two dimensional histograms denoting Ramachandran (columns 1,2) and Janin (columns 3,4) plots for (top row) glycosylated mini mucins and (bottom row) unglycosylated mini mucins. Within the sets of Ramachandran and Janin plots, the first column is all Φ/Ψ dihedral distributions for non-glycosite residues, the second column is all Φ/Ψ dihedral distributions for glycosite residues. 2D-hexagonal histogram bins are colored according to population density on a kcal/mol scale. A molecular graphic is shown on the right to demonstrate which torsions are considered (mucin protein backbone atoms in red; carbon, nitrogen, and oxygen, atoms are highlighted in black, blue, and red spheres, respectively, O-glycan atoms highlighted in yellow, neighboring glycan atoms shown in light grey). (B) Network graphs highlighting the average number of glycan-glycan contacts (line color) and standard deviation in this average (line width) for Mini1 (left) and Mini2 (right). Charged glycans are represented with red circles, neutral glycans are represented with grey circles. Inset images show examples of contacts between “far” neighbor glycans, and inset bar graphs depict the average number of glycan-glycan contacts between each pair-type. (C) Left and middle panels: First and second water solvation shell counts for protein (pink and red) and glycans (cyan and blue). Right panel: Solvation shells considered in analysis: glycans (grey licorice), protein (black licorice), first and second solvation shells shown in shades of blue and red.

Furthermore, with the addition of an O-glycan on a Ser or Thr residues, glycosylated Ser and Thr now have a non-trivial χ2 torsional potential and thus Janin plots for such glycosites in glycosylated mini mucin simulations are drastically different compared to those same residues in unglycosylated mini mucin simulations. The addition of the true χ2 results in a dramatic reduction of degrees of freedom. We compare these Janin plots to non-glycosylated residues (i.e., not Ser or Thr) with true χ2 dihedrals from our Mini mucin simulations, specifically a His and Leu in each mini mucin. As can be seen, not only do glycosite residues demonstrate dramatically different Janin plots between glycosylated and unglycosylated mini mucin simulations (as to be expected as Ser and Thr do not have true χ2s when unglycosylated) but they also have dramatically different Janin distributions relative to His and Leu in glycosylated and unglycosylated mucin simulations. These factors demonstrate the reduced degrees of freedom accessible to mucin protein backbones due to O-glycosylation. All our Ramachandran and Janin plots agree remarkably with NMR and early restrained minimization modeling data from Coltart et al^85^ who demonstrated that modification by the first O-GalNAc residue was sufficient to impose these restraints to protein backbone conformational torsions. We also calculated the extent of glycan-glycan contacts (defined as the number of residues from one glycan within 5Å of another glycan) within all glycosylated mini mucin simulations (**Figure 6B**). Contact networks show the average number of contacts as well as standard deviation in this average between all glycans in each mini mucin. Overall, we see that most glycans do contact most other glycans within each mini mucin. However, we see that the strongest degrees of glycan-glycan contacts are those within the “nearest neighbor” range, inset bar graphs in **Figure 6B**, i.e., 60% of all glycan-glycan contacts within each mini mucin system are within the first 3 nearest neighbors (see supporting information **Figure S16** for full integration plots). We hypothesize that the nearest neighbor glycan-glycan contacts underlie the increasing persistence length exhibited by the glycosylated mucin systems. We also calculated the extent of glycan-glycan contacts (defined as the number of residues from one glycan within 5Å of another glycan) within all glycosylated mini mucin simulations (**Figure 6B**). Contact networks show the average number of contacts as well as standard deviation in this average between all glycans in each mini mucin. Overall, we see that most glycans do contact most other glycans within each mini mucin. However, we see that the strongest degrees of glycan-glycan contacts are those within the “nearest neighbor” range, inset bar graphs in **Figure 6B**, i.e., 60% of all glycan-glycan contacts within each mini mucin system are within the first 3 nearest neighbors (see **Figure S12** for full integration plots). We hypothesize that the nearest neighbor glycan-glycan contacts underlie the increasing persistence length exhibited by the glycosylated mucin systems.

### Solvation promotes extended mucin conformations

Kramer et al.,^84^ hypothesized the solvation shell of O-glycans may contribute to steric hinderance that enforces extended mucin protein backbones. Following this line of reasoning, we sought to characterize the density of solvation shells surrounding glycosylated mini mucins and unglycosylated mini mucins during simulation. To do so, for each frame in simulation, we calculated the number of water molecules within the first and second solvation shells (3.4 and 5 Å), respectively, of glycan segments or protein residues (**Figure 5C**, supporting information **Figure S17**). The average number of water molecules in the first and second solvation shells of protein residues in unglycosylated mini systems (protein-only) was 13 ± 4 and 32 ± 9, respectively, with variability in this average stemming largely from residue size (i.e., per residue values have much lower variability). By comparison, the number of water molecules within the first and second solvation shells of mucin residues (i.e., considering O-glycans) was 65 ± 40 and 123 ± 72 respectively (the average number of water molecules within first and second solvation shells of protein residues within glycosylated mucins was 8 ± 4 and 19 ± 9, respectively). Thus, O-glycans result in an order of magnitude increase in the number of water molecules within the first solvation shell in the glycosylated mucin.

We additionally sought to characterize dynamics of the water molecules around the mini mucin models in glycosylated and unglycosylated states. Water molecules within the first solvation shell of glycosylated mucins have a residence time of 70 ± 3 ps while water molecules within the first solvation shell of unglycosylated mini mucins have a residence time of 57 ± 4 ps (**Figure S18A);** thus, water molecules around glycosylated mucins remain bound for on average ∼10 ps longer. Furthermore, mean square displacement (MSD) calculations reveal that water molecules within the first solvation shell of glycosylated mucins move faster on average than water molecules within the first solvation shell of unglycosylated mucin models (1.5 ± 0.1 x10^-7^ m^2^/s and 1.1 ± 0.2 x10^-7^ m^2^/s, respectively **Figure S18B**). Interestingly, water molecules within the first solvation shell of glycosylated mucins move on average as fast (1.5 ± 0.1 x10^-7^ m^2^/s) as water molecules in bulk conditions (1.50 ± 0.02 x10^-7^ m^2^/s) while water molecules in the first solvation shell of unglycosylated mucins move more slowly (1.1 ± 0.2 x10^-7^ m^2^/s) than waters in the bulk (**Figure S18B**). We hypothesize that difference in water molecule behavior within first solvation shells of glycosylated and unglycosylated mucin and bulk conditions may be due to the number of available hydrogen bonding partners supported by O-glycans. Mucin O-glycans each have many hydroxyl and charged moieties that can each donate or accept hydrogen bonds and thus water molecules can exchange, translate, and rotate, more readily without significantly sacrificing hydrogen bonding partners. Thus, extended mucin protein backbones allow for maximization of such hydrogen bonding partners with water molecules. However, unglycosylated mucins are relatively more hydrophobic in nature; each unglycosylated Ser and Thr can only accept one and donate one hydrogen bond, but the remaining residues are largely hydrophobic (Gly, Pro, Leu, and Val). Thus, for the case of the unglycosylated mucins, we hypothesize the hydrophobic effect contributes to protein collapse, in turn minimizing solvent accessible surface area.

### Mega mucin model captures flexibility demonstrated from smaller simulations while recapitulating experiment

Finally, we sought to determine whether our Mega mucin model simulations agree qualitatively with experimentally characterized images of physiological mucins. Towards this aim, Anggara et al. graciously provided us with twenty scanning tunneling microscopy images of soft-landing electrospray ion beam deposited MUC1 tandem repeat fragment (6.5 tandem repeats around 130 amino acids) produced in glycoengineered cells.^54,59^ Each STM image is an image of a single MUC1 molecule. Although MUC1 and MUC5B are different in both PTS domain tandem protein backbone repeats and specific O-glycoprofiles, the qualitative similarities between our model (**Figure 7B**) and these experimentally acquired images available in **Figure 7A** and **7C** are too remarkable to ignore. To normalize for the size differences of our mega mucin and Anggara et al.’s MUC1 images, we measured end-to-end distances and divided these results by the number of amino acid residues present in each construct (224 and 148, respectively, Supporting Information **Figure S19**). A detailed analysis description and MUC1 images can be found in SI. Although there does not seem to be significant agreement between the normalized average height per residue values for MUC1 vs our mega mucin model (Supporting Information **Figure S20**), this can be largely attributed to the technique of macromolecule deposition for STM imaging wherein mucins are deposited on a surface and stretched to achieve maximum clarity on O-glycans.

**Figure 7:**
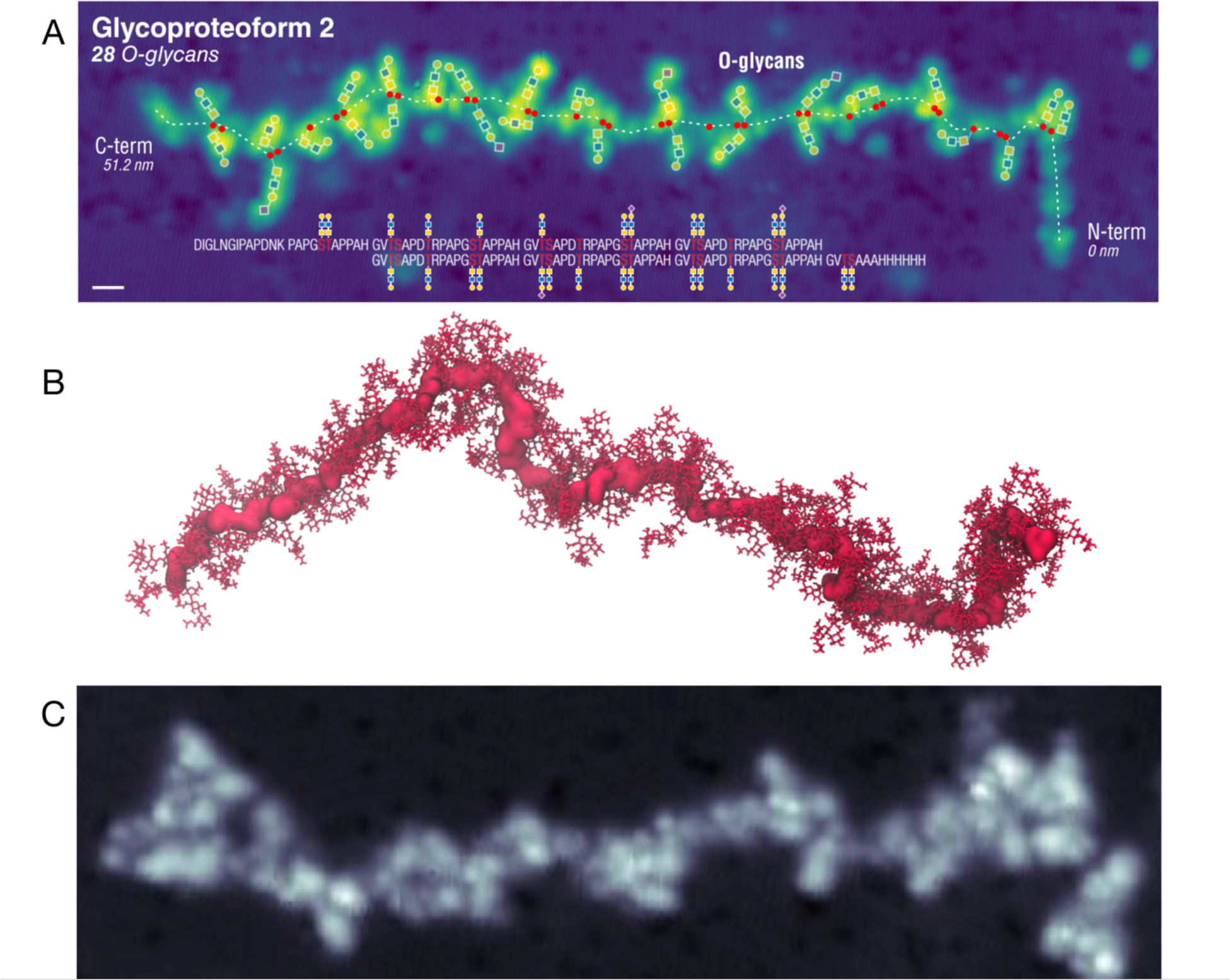
Direct comparisons of computational Mega mucin to experimentally derived 2d images of MUC1. (A) Scanning tunneling microscopy image of soft-landing electrospray ion beam deposited MUC1, annotated by Anggara et al.^59^ (B) 148 amino acid section of a representative frame mega mucin simulation rendered with protein as a red surface and glycans as red licorice. (C) Another representative scanning tunneling microscopy image from Anggara et al demonstrating qualitative similarity to (B). Collectively, our results suggest that the bottlebrush phenomenon is derived from: (1) reduced torsional degrees of freedom along the protein backbone due to glycan induced steric hindrance, (2) nearest neighbor glycan contacts which amplify reduction in torsional potentials, and (3) the extension of the solvation shell which further amplifies glycan steric hindrance as well as maximization of hydrogen bonding potential to solvent.

## Conclusions

From our miniature mucin simulations, we have delineated several key degrees of freedom that impart physical structure to otherwise intrinsically disordered mucin protein backbones: (1) decreased backbone torsional degrees of freedom, (2) strong nearest-neighbor contacts between O-glycans in densely glycosylated mucins, and (3) dramatically increased first and second solvation shells from the introduction of hydrophilic glycans. Furthermore, the internally consistent persistence length calculated for Mini and Mega mucin models provides great confidence in using such mini-mucin models to accurately and inexpensively approximate mucin biochemical properties and mechanics, for example, when analyzing overall interaction profiles between mucin and other proteins. In addition to validating our mini mucin models, our Mega mucin model provides a striking qualitative comparison to experimentally derived images of MUC1. Further, the development of small (∼200 amino acid) mucin reporter constructs with tailored O-glycan positions, as done by Nason et al,^19,54^ allows for potential collaborative exploration via experiment and simulation with reasonable overlap in mucin sequence and structure. Similarly, expansion of glycomics methods, as done by development of mucinases and LC-MS/MS techniques, will allow for increasing accuracy in O-glycan structures which can then feedback into the workflow for increased predictive power in mucin modeling.^56,60^ Finally, development of novel visualization techniques for mucins will dramatically strengthen relationships between mucin simulation and experiment.^59^

Due to their many roles, dysregulation and altered morphology of mucins are implicated in many diseases including diabetes mellitus, many cancers, and cystic fibrosis. As such, understanding the basic nature of mucins and their O-glycans in imparting structure to the mucosal layer, is vital to understanding disease pathogenesis and progression. Future investigations into the molecular mechanisms of these diseases may require in-depth, atomic-scale, structural understanding of mucin glycoproteins. We have demonstrated the construction and simulation of densely O-glycosylated mucin domains, and how these simulations can be leveraged to investigate the biochemical/biophysical nature of mucin domain glycoproteins. The simulations presented herein represent a starting point through which novel structure/function hypotheses can be explored in this domain.

## Acknowledgements

We thank Kelvin Anggara and Rebecca Miller for their generosity in allowing us to use their STM images of MUC1, and Thapakorn Jaroentomeechai and Yoshiki Narimatsu who designed and produced the MUC1 mucin reporter used in that work. We also thank Elisa Fadda, Ronit Freeman, Stacy Malaker, Kelvin Anggara, Rebecca Miller, and Henrik Clausen (and their labs) for invaluable discussions and insights into mucin glycobiology. We acknowledge the San Diego Supercomputing Center (SDSC) for providing HPC resources that have contributed to the research results reported within this paper.

## Abbreviations

SNFG: Symbol Nomenclature for Glycans
MD simulations: Molecular Dynamics Simulations
MUC5B: canonical mucin 5B
PL: Persistence Length
Rg: Radius of Gyration
RMSF: Root Mean Square Fluctuation
RMSD: Root Mean Square Deviation
RLS: Relative Linearity Score
Glycosylated Mini1: Miniature Mucin Model #1, simulations performed with glycosylation
Glycosylated Mini2: Miniature Mucin Model #2, simulations performed with glycosylation
Unglycosylated Mini1: Miniature Mucin Model #1, simulations performed of just protein backbone
Unglycosylated Mini2: Miniature Mucin Model #2, simulations performed of just protein backbone
Glycosylated Mega: Large Mucin Model, simulations performed with glycosylation
Unglycosylated Mega: Large Mucin Model, simulations performed of just protein backbone
STM: scanning tunneling microscopy

## Data Sharing

Trajectories, structures, simulation scripts, analysis scripts, and data files will be shared with this work through a link hosted on the UC San Diego library website.

