## Supporting Information for "Breaking Down the Bottlebrush: Atomically-Detailed Structural Dynamics of Mucins"

#### Table of Contents:

1. Methods:
 1.1 Miniature Glycosylated and Unglycosylated Simulations
 1.2 Mega Glycosylated and Unglycosylated Simulations
 1.3 Analysis Methods
  1.3.1 Principal Component Analysis and Kmeans Clustering
  1.3.2 Persistence Length Calculations
  1.3.3 Relative end-to-end distances (computational)
  1.3.4 Relative Radius of Gyration
  1.3.5 Relative compactness
  1.3.6 Curvature per $C_\alpha$
  1.3.7 $C_\alpha$ Pinch Points
  1.3.8 Cluster Linearity Score Calculations
  1.3.9 Protein Root Mean Square Fluctuations
  1.3.10 Glycan Root Mean Square Fluctuations
  1.3.11 Ramachandran and Janin Plots
  1.3.12 Glycan-Glycan Contacts and Network Generations
  1.3.13 Solvation Shells
  1.3.14.1 Length Per Residue (Computational)
  1.3.14.2 Length Per Residue (Experimental)
  1.3.15.1 Bending Angle Calculations (Computational)
  1.3.15.2 Bending Angle Calculations (Experimental)

2. Results and Discussion:
 2.1 Figures:
  **Figure S1:** Construction of Unglycosylated mini mucin systems.
  **Figure S2:** Glycosylated mini mucin clustering statistics.
  **Figure S3:** Per-cluster descriptions from the 11 clusters collected from glycosylated mini mucin simulations.
  **Figure S4:** General properties from glycosylated mini1 and mini2 simulations.
  **Figure S5:** Unglycosylated mini mucin clustering statistics.

**Figure S6:** Per-cluster descriptions from the 11 clusters collected from Unglycosylated mini mucin simulations.
**Figure S7:** General properties from Unglycosylated mini1 and mini2 simulations.
**Figure S8:** Pinch points per residue for glycosylated (A) Mini1 and (B) Mini2 simulations.
**Figure S9:** Pinch points per residue for Unglycosylated (A) Mini1 and (B) Mini2 simulations.
**Figure S10:** Relative Rg per glycan from glycosylated (A) Mini1 and (B) Mini2 simulations.
**Figure S11** Glycosylated mega mucin clustering statistics.
**Figure S12** Absolute end-to-end distances of glycosylated mega mucin system color coded by cluster identify.
**Figure S13:** Principal component analysis of concatenated three replicas of glycosylated mega mucin system and one replica of unglycosylated mega mucin system.
**Figure S14**: RMSFs calculated for glycosylated and unglycosylated mega mucin system.
**Figure S15:** Φ/Ψ/χ1/χ2 1D probability densities.
**Figure S16:** Full-size bar graphs and integration plots demonstrating the number of contacts between glycans and their neighbors.
**Figure S17:** Solvation shells per glycan.
**Figure S18:** Water behavior around glycosylated and unglycosylated mini mucin models.
**Figure S19:** Scanning tunneling microscopy image of soft-landing electrospray ion beam deposited MUC-1 graciously provided by Anggara et al.
**Figure S20:** Normalized measurement of height per residue in angstroms for Mega mucin and experimental STM MUC-1 images.

3. References

### 1. Methods:

***1.0 Step-by-step guide to constructing Mini1 and Mini2 models:*** Protein sequence acquisition: Hughes et al[5] published a logogram detailing the consensus sequence repeats most likely seen in MUC5B PTS domains. From their logogram, we selected a 26 amino acid sequence using all the most likely amino acids found at each position (Hughes et al.[4] Figure 3). This peptide became the backbone for Mini1. Additionally, Hughes et al [4] also published a sequence consistent with consensus sequence repeat patterns most likely seen in MUC5B PTS domains (Hughes et al.[4] Figure 6); we used that sequence to construct a 30 amino acid polypeptide, this peptide then served as the backbone for Mini2. We used Rosetta's BuildPeptide tool[44] to build peptide structures of Mini1 and Mini2 sequences. Identification of stereo-nonspecific Mass Spectromery Data for glycan structures: Extensive mass spectrometry data from Thomsson et al[6] isolates various O-glycan structures from respiratory mucosal samples. However, these data did not provide stereochemistry nor connectivity information. Collection of stereospecific glycan structures from the GlyConnect Expasy Database: Cross comparison of the list of stereo-non-specific Thomsson et al[6] data to the extensive list of O-glycans isolated in pulmonary mucosa from the GlyConnect Database, which includes stereochemistry and connectivity information.[45] We selected O-glycans from the GlyConnect Expasy database that corresponded to glycans with masses comparable to molecular ions listed in Thomsson et al's[6] Table 1. Additionally, we selected a variety of charged and neutral glycans, and we took care to select more Core-1 and Core-2 glycotypes than Core-3 and Core-4 glycotypes as the former are slightly more common. This set of O-glycans served as our "in-house" library of glycans. Construction of mucin models: CHARMM-GUI[46] Glycan Modeller[47] was used to add O-glycan structures from steps 2+3 to our Mini1 and Mini2 protein sequences identified in step 1. Here, we considered all Ser and Thr residues on Mini1+2 sequences to be potential O-glycosites, and thus modified them with selected O-glycans (except Thr26 on Mini1, which was left unglycosylated). We then "pseudo-randomized" the placement of selected O-glycan structures onto their assigned mucin models; charge densities were distributed along the mucin model to avoid clustering of charged glycans within mucins. The resulting O-glycoprotein maps for Mini1 and Mini2 can be seen in **Figure 3A** and see **Table 1** for proportions of Core types and charged groups within Mini1 and Mini2 models.

***1.1 Miniature Glycosylated and Unglycosylated Simulations:*** All systems (including Unglycosylated miniature systems **Figure S1**) were simulated with NAMD3[1,2] using SDSC resources and parameters described by CHARMM36m all-atom force-fields.[3–5] For all simulations: Periodic Boundary Conditions were used, Particle Mesh Ewald was used to calculate long-range Coulombic potentials, Lennard-Jones potentials were calculated according to a 10 Å – 12 Å cutoff and switching scheme, pairwise list generation was cutoff at 15.5 Å. Langevin thermostat and piston were used to maintain temperature at the following specified temperatures and at pressure of 1 atmosphere. *Minimization:* To relax all conformations and to alleviate close contacts resultant from O-glycosylation, all solvated and neutralized structures were subjected to 10,000 steps of conjugate gradient minimization with no atomic restraints or constraints. *Heating:* Systems were then heated from 10 K to 310 K, increasing temperature by 25 K every 10,080 ps, timestep 2 fs, for a total of 240 ps of heating. Once at 310 K, an additional 770 ps of simulation was conducted, jumpstarting a short equilibration. *Equilibration:* Systems were then subjected to NpT equilibration at 310 K for 0.5 ns, 2 fs timestep (useFlexibleCell option "on", wrapWater option "on", wrapAll option "on"). Systems were then subjected to an additional 55 ns NVT equilibration with 2 fs timestep with fixed box dimensions (useFlexibleCell option "off"). *Production*: Finally, 550 ns of NVT production runs were conducted at 310 K, timestep 2 fs. All data presented in the Results and Discussion section for glycosylated and Unglycosylated mini mucin systems is derived from these 550 ns production runs.

***1.2 Mega Glycosylated and Unglycosylated Simulations:*** Simulations for mega glycosylated system were performed according to the methodology established in 1.1, but with a few key differences: 1) system was subjected to an additional 360 ns of equilibration in order to acquire a structure that was in a state of pseudo-equilibration 2) a frame was picked at 360 ns after analysis was done to determine the system was no longer compressing from its artificial outstretched state, i.e. no longer rapidly decreasing in end-to-end distance, and 3) as an effort to perform adaptive sampling on an otherwise prohibitively large system, simulations were launched from the extracted pseudo length equilibrated frame at reinitialized velocities for 210 ns, 283 ns, and 264 ns for replica 1, replica 2, and replica 3, respectively.

***1.3 Analysis Methods:*** Unless otherwise stated, the following analyses were performed with MDAnalysis.[6,7] All analysis scripts are shared along with shared trajectories and resultant data files along with this Supporting Information.

1.3.1 Principal Component Analysis and Kmeans Clustering: We sought to perform shared states clustering within all glycosylated and within all Unglycosylated simulations. That is to say, we sought to, for example, find a way to concatenate all simulation frames from Mini1 and Mini2 glycosylated simulations, align those trajectories and cluster those conformations together. Doing so would allow us to compare the morphology of clusters adopted by Mini1 versus Mini2. Similarly, if we could perform similar shared state clustering for Unglycosylated mucin simulations then we could similarly identify which cluster morphologies were populated for Mini1 versus Mini2 peptides. However, given the fact that Mini1 and Mini2 were heterogeneously constructed for the purposes of more accurately representing heterogeneity observed in the mucosal layer, to concatenate the trajectories we had to first do the following: (1) Strip all trajectories to their backbone atoms only, removing atoms of all waters, ions, side chains, and glycans, (2) truncate mini2 trajectories by removing terminal residues so that both Mini1 and Mini2 peptides had the same length (we removed residues 1,2,29, and 30), (3) rename all residues to Glycine, and (4) write psf files consistent with these new trajectories. Following this harsh stripping procedure, for each trajectory (Mini1 vs. Mini2, Unglycosylated vs. glycosylated, replicas 1, 2, or 3) we were left with backbone conformations only, which was ideal for this intended purpose. For glycosylated simulations, stripped trajectories that corresponded to simulations performed with glycans were all concatenated with the same topological information (same psf) using MDAnalysis. The concatenated trajectory was then aligned (using the AlignTraj class from MDAnalysis's align module[6–9]) to a reference minimized outstretched structure that was also stripped, using $C_\alpha$ atoms as the alignment selection. Following alignment, MDAanalysis's built-in principal component analysis tool (PCA class) was used to calculate the principal components that explained 95% of variance in $C_\alpha$ atomic positions in the concatenated and aligned trajectory. Following completion of PCA analysis, we plotted variance and fraction of cumulated variance for all identified PCs, **Figure S2.** PSF_pca identified 78 PCs explained 95% of variance in the case of glycosylated simulations, **Figure S2.** The top 3 PCs were selected for clustering as those were able to sufficiently describe over 60% of variance (70%) in $C_\alpha$ positions, **Figure S2**. We plotted all simulation frames in a 3D scatter plot according to their calculated PCs, we colored those points by frame number, **Figure S2.** We then clustered all frames according to the top 3 PCs, using scikit-learns's kmeans clustering module.[10] We chose the optimal number of clusters to be that which satisfied both the "elbow test", i.e. was past the inflection point in the sum of squares error per cluster, and the silhouette coefficient test, i.e., produced a silhouette coefficient that was greater than 1 standard deviation above the average silhouette coefficient taken across all clusters, **Figure S2.** For glycosylated simulations, the optimal number of clusters was determined to be 11. For each cluster, we identified the most representative structure as that which having the lowest RMSD relative to the cluster center in PC space. We also calculated distributions of several properties for all frames in each cluster including relative end-to-end distances, relative radii of gyration, and relative compactness, see below for how these values are calculated. From these values, we derived a "linearity score" (see below) describing the mucin peptide backbone conformations. The median linearity score per cluster was used to qualitatively rank each cluster in terms of "least" to "most" "linear". We then plotted all resultant descriptive values on a per-cluster basis, **Figure S3**. The above clustering procedure was then performed for the stripped Unglycosylated mini mucin trajectories as well. See below table for comparative list of PCA and kmeans clustering statistical values:

| | Glyco Mini Clustering | Deglyco Mini Clustering |
|---|---|---|
| Number of Frames Clustered | 660,000 | 660,000 |
| #PCs to explain 100% var | 78 | 78 |
| #PCs to explain 95% var | 11 | 11 |

| | | |
|---|---|---|
| #PCs to explain 60% var | 3 | 3 |
| Optimal #Clusters | 11 | 11 |

1.3.2 Persistence Length Calculations: Persistence length was calculated for each model in each simulation, in each replica, using MDAnalysis's persistence length class in the polymer module.[6,7] The polymer of interest was selected to be all the $C_\alpha$ atoms in each mucin backbone. We calculated the average (and standard deviation) of all persistence lengths over each mucin modeled, i.e., Mini1 versus Mini2 in Unglycosylated versus glycosylated states, and Mega mucin in Unglycosylated and glycosylated states. We saw striking agreement between persistence lengths calculated for Mini1 and Mini2 **(**see **Figure S4),** thus we deemed it appropriate in the main text to report the average persistence length for Unglycosylated mini simulations versus glycosylated mini simulations versus glycosylated mega simulations.

1.3.3.1 Relative end-to-end distances (computational): Relative end-to-end distance was calculated for every frame in each simulation by first calculating the end-to-end distance along the protein backbone of each given structure (i.e., Mini1 or Mini2 in Glycosylated states, Mini1 or Mini2 in Unglycosylated states, or Mega mucin in glycosylated or Unglycosylated states) by calculating the distance between the first $C_\alpha$ atom and the last $C_\alpha$ atom in the minimized/outstretched state. This was referred to as the "reference end-to-end distance". Then, for every frame in the simulation, the ratio between the instantaneous/framewise end-to-end distance (distance between the first $C_\alpha$ atom and the last $C_\alpha$ atom) and the reference end-to-end distance was calculated. Thus, a value approaching 1.0 for a relative end-to-end distance indicates the length of the protein backbone in that given frame is close to its value in the minimized state. Similarly, a relative end-to-end distance approaching 0.0 indicates the distance between the first and last $C_\alpha$ atom is highly compacted relative to the minimized/outstretched state, and a value close to 0.5 indicates the distance between the first and last Cα atom is around half of the distance seen in the minimized/outstretched state.

1.3.4 Relative Radius of Gyration: Relative radius of gyration (relative Rg) was calculated for every frame in each simulation by first calculating the Rg of each structure (i.e., Mini1 or Mini2 in Glycosylated states, Mini1 or Mini2 in Unglycosylated states) in the minimized/outstretched stated. This was referred to as the "reference Rg". Then, for every frame in the simulation, the ratio between the instantaneous/framewise Rg and the reference Rg was calculated. Thus, a value approaching 1.0 for relative Rg indicates the framewise Rg is close to that seen in the minimized state. Similarly, a relative Rg approaching 0.0 indicates the framewise Rg is much less than that seen in the minimized/outstretched state. Relative Rgs for glycosylated and Unglycosylated mini mucins were calculated by selecting "all" atoms in the respective systems. However, relative Rg's for glycans were calculated by selecting each specific glycan.

1.3.5 Relative compactness: Relative compactness was calculated for every frame in each simulation by first calculating the "compactness" of all $C_\alpha$'s relative to the protein backbone center of mass of each structure (i.e., Mini1 or Mini2 in Glycosylated states, Mini1 or Mini2 in Unglycosylated states) in the minimized/outstretched stated. This was referred to as the "reference compactness". The exact equation used to calculate $C_\alpha$ compactness was as follows:

Compactness = np.sqrt((np.sum(cas.positions – peptide.center_of_geometry())**2)

Where peptide.center_of_geometry() was taken as the center of mass of all $C_\alpha$s along the protein backbone. Then, for every frame in the simulation, the ratio between the instantaneous/framewise compactness and the reference compactness was calculated. Thus, a value approaching 1.0 for relative compactness indicates the framewise $C_\alpha$ compactness is close to that seen in the minimized state. Similarly, approaching 0.0 indicates the framewise $C_\alpha$ compactness is less than that seen in the minimized/outstretched state.

1.3.6 Curvature per $C_\alpha$: Curvature per $C_\alpha$ atom per simulation frame for all systems (i.e., Mini1 or Mini2 in Glycosylated states, Mini1 or Mini2 in Unglycosylated states) was calculated using the following equation for curvature:

Curvature = np.sqrt (( d2z*dy – d2y*dz)**2 + (d2x*dz – d2z*dx)**2 + (d2y*dx – d2x*dy)**2/(dx**2 + dy**2 + dz**2)**1.5

Where dx, dy, dz, d2z, d2y, d2z, were calculated as x,y,z coordinates first and second derivatives around each Cα (estimated by first and second derivatives between $C_{\alpha-1}$ and $C_{\alpha+1}$, except for end point $C_\alpha$s which were estimated as $C_\alpha$ to $C_{\alpha+1}$ or $C_{\alpha-1}$ to $C_\alpha$ as necessarily).

1.3.7 $C_\alpha$ Pinch Points: Following completion of MD simulations, we observed that several frames and several representative cluster centers demonstrated "pinched" protein backbones wherein the backbone changed "direction" significantly around 1-2 amino acids. This can be seen prominently in the representative cluster structures for glycosylated Cluster 2, and several of the representative structures for the Unglycosylated mucin clusters. To describe these "pinch points" along each simulation we sought to calculate the angle around each $C_\alpha$ per simulation frame. To calculate the angles around each $C_\alpha$ atom, for each simulation we calculated the angle formed between each $C_{\alpha-1}$, $C_\alpha$, and $C_{\alpha+1}$, for each frame in the simulation, excluding the first and last $C_\alpha$s. alpha-helices are characterized by a 360° rotation in 3.6 amino acids, thus the angle formed between $C_{\alpha-1}$, $C_\alpha$, and $C_{\alpha+1}$ for amino acids in an alpha-helix corresponds to ~100°. Thus, we considered any angle formed between $C_{\alpha-1}$, $C_\alpha$, and $C_{\alpha+1}$ with a value less than 100° to be a "pinched" angle, as this would represent a $C_\alpha$ atom with a greater "turn" than that seen in alpha-helices. We then the number of these "pinched $C_\alpha$s" per frame and for the Unglycosylated and glycosylated mini mucins we calculated the average number of "pinched $C_\alpha$s" per frame.

1.3.8 Cluster Linearity Score Calculations: To qualitatively describe and compare clusters from mini mucin clustering, and to rank and compare clusters between glycosylated and Unglycosylated mini mucin clustering conditions, we derived a "cluster linearity score". Per simulation frame we calculated a framewise linearity score which was taken as:

Framewise linearity score = relative end-to-end distance * relative Rg * relative compactness

The cluster score was then taken as the median of all framewise linearity scores calculated from the set of linearity scores from all frames in that cluster:

Cluster linearity score = median (Framewise linearity scores per cluster)

1.3.9 Protein Root Mean Square Fluctuations: Root mean square fluctuations (RMSFs) were calculated for Mini1 and Mini2 glycosylated and Unglycosylated systems using the MDAnalysis

RMSF functionality. Before RMSF was calculated for all systems, each trajectory was aligned to a reference starting frame: the first replica's minimized/outstretched structure for the Mini1/Mini2/glycosylated/Unglycosylated simulation as appropriate. RMSF was then calculated for each $C_\alpha$ in each system.

1.3.10 Glycan Root Mean Square Fluctuations: Glycan RMSF was performed by first, per glycan, aligning the target glycosylated trajectory to the minimized starting structure using the heavy atoms of the protein glycosite (i.e., the Ser or Thr modified by the target O-glycan) and the glycan segments's first residue as the alignment selection. Then, we used MDAnalysis RMSF calculator to calculate the RMSF for all $C_1$ atoms in the glycan segment, which we took as an analog to $C_\alpha$s in protein segments. To report a singular "Glycan RMSF" per glycan, we then calculated the average and standard deviation of these "per C1" RMSFs over the whole glycan. This is also why the standard deviations for glycan RMSFs are broader than the protein RMSFs: since we chose to align the glycans only by their protein glycosite and their first residue, terminal glycan residues have higher RMSFs than the first residue. We also calculated glycan RMSFs by first aligning trajectories to the first frame using the whole glycan position as the target for alignment, but we felt these results gave artificially low values due reducing too many degrees of freedom and completely canceling protein motions. We felt alignment by protein glycosite, and first glycan segment's residue provided the most holistic and faithful demonstration of glycan flexibility.

1.3.11 Ramachandran and Janin Plots: For glycosylated and Unglycosylated mini mucin simulations, Ramachandran plots ($\Phi/\Psi$ distributions) were calculated using MDAnalysis's Ramachandran class in the "dihedrals" module, excluding terminal residues. Data was saved in the form of pandas DataFrames where specific residues could be pulled for binning and plotting, i.e., glycosite vs non-glycosite data was parsed post-analysis. Similarly, Janin plots ($\chi1/\chi2$) for glycosylated and Unglycosylated mini mucin simulations were calculated using the Janin class in the "dihedrals" module. The Janin class automatically pulls amino acids from a selection for which there are non-trivial $\chi1$ and $\chi2$ angles, meaning it skips amino acids without $\chi1$ or $\chi2$ dihedrals which are completely defined by heavy (non-hydrogen) atoms. In the case of our Mini1 and Mini2 sequences, the only non-glycosite residues that satisfy this requirement are thus H15 and L21 in Mini1 and L2 and H27 in Mini2. However, in the case of Ser and Thr, when these residues are modified by O-glycans, they then have a $\chi2$ angle that is not trivial. Therefore, we used MDAnalysis distances library, calc_dihedrals function to calculate these $\chi1$ and $\chi2$ dihedrals as the Janin class could not be coaxed to select $\chi2$s for O-glycans and their glycosites. We then used the calc_dihedrals to also calculate dihedrals for Ser and Thr residues in the Unglycosylated mini mucin simulations, although we acknowledge that these residues have a somewhat trivial "$\chi2$" torsional potential.

1.3.12 Glycan-Glycan Contacts and Network Generations: For all glycosylated miniature simulations, we sought to tabulate the number of glycan-glycan contacts to estimate the degree to which glycan contacts facilitated within-mucin communication which could impact persistence length. A glycan-glycan contact was calculated as when a glycan residue in one glycan segment was found to be within 5 Angstroms of a glycan residue on another glycan segment (i.e., glycan contacts within glycan segments were not counted in this analysis). Per frame, the number of glycan contacts made by each glycan was tabulated in DataFrames files. Per glycan, we then calculated the average number of (and standard deviation of) contacts between each glycan and every other glycan. We then used the python NetworkX module to display the average number of contacts and standard deviation in that average between each glycan-glycan pair in Mini1 and Mini2 glycosylated simulations. We then calculated the average number and standard deviations of contacts between each "type" of glycan-glycan pair, i.e., whether those glycans were next to each other, one glycan away from each other, two glycans away from each other, and so on. We then calculated, for pairs of neutral glycans, neutral and charged glycans, and charged glycans, the average number of contacts between each glycan chemical pair type.

1.3.13 Solvation Shells: We calculated the number of water molecules in the first and second solvation shells by using MDAnalysis selection commands to iteratively select for and count the number of water molecules within 3.4 and 5.0 Å, respectively, of protein residues or glycan segments per frame. We then calculated the average number of, and standard deviation of, these water molecules in each region. We did not account for double counting: meaning, for example, some water molecules could be found to be both in the first solvation shell of a protein residue and in the first or second solvation shell of a glycan residue, however, at this time we don't suspect this would lead to unreasonable degrees of error due to the dramatic differences in numbers of water molecules accounted for around glycan segments.

1.3.14.2 Length Per Residue (computational): Relative end-to-end distance was calculated for every STM image provided by Clausen et al. by first calculating the end-to-end distance along the protein backbone of MUC-1 by approximating the distance between the first $C_\alpha$ atom and the last $C_\alpha$ atom in the minimized/outstretched state. This was referred to as the "reference end-to-end distance". Then, for every frame in the simulation, the ratio between the instantaneous/framewise end-to-end distance (distance between the first $C_\alpha$ atom and the last $C_\alpha$ atom) and the reference end-to-end distance was calculated. Thus, a value approaching 1.0 for a relative end-to-end distance indicates the length of the protein backbone in that given frame is close to its value in the minimized state. Similarly, a relative end-to-end distance approaching 0.0 indicates the distance between the first and last $C_\alpha$ atom is highly compacted relative to the minimized/outstretched state, and a value close to 0.5 indicates the distance between the first and last Cα atom is around half of the distance seen in the minimized/outstretched state.

1.3.15 Water Residence Time Calculations: Water residence times were calculated using the survival probability class in MDAnalysis's WaterDynamics module.[11–17] For every 10,000$^{th}$ frame in our 110,000 frame simulations (i.e., every 50ns out of the 550 ns simulations) we selected all water molecules within the first solvation shell (<3.4 Å) of a given selection (e.g., within the first solvation shell of all mucin atoms, all glycan atoms, or all protein atoms) and we calculated the survival probability of those waters to remain within that solvation shell for 500 ps. We then estimated the residence time by taking the inflection point in the smoothed survival probability curve: i.e., we found the time frame over which water molecules within the first solvation shell of a given selection lost significant probability for remaining in that solvation shell. This procedure was repeated for all See Shared Files for this calculation function.

1.3.16 Water Mean Square Displacement Calculations: Water mean square displacements (MSDs) were calculated using the mean square displacement class within MDAnalysis's WaterDynamics module.[11–17] For every 10,000$^{th}$ frame in our 110,000 frame simulations (i.e., every 50 ns out of 550 ns simulations) we selected all water molecules within the first solvation shell (<3.4 Å) of a given selection (e.g., within the first solvation shell of all mucin atoms, all glycan atoms, or all protein atoms) and we calculated the mean square displacement of waters within that solvation shell for the next 500 ps. To determine the mean square displacement for water molecules in bulk water conditions, we also performed three replicas of 550 ns production simulation (simulated identically as described in Supporting Information Methods Section 1.1) of a 150 mM NaCl water box (see table below for bulk water system details). From these water box simulations, every 50 ns we then also calculated the MSD of all water molecules for the next 500 ps. These “bulk water” results serve as a standard for comparison for waters in glycosylated and unglycosylated mini mucin simulations.

| | Bulk Water |
|---|---|
| Total #atoms | 50,924 |
| #protein atoms/residues | 0 |
| Protein Backbone Length (Å) | -- |
| #glycan atoms/ #glycans | 0 |
| #water atoms/residues | 52,407 / 17,469 |
| #Na/#Cl ions | 49 / 49 |
| System dimensions (AxAxA) | 63.9 x 68.7 x 124.99 |
| Total Sampling | 3 x 550 ns |

#### 2. Results and Discussion:

##### 2.1 Figures:

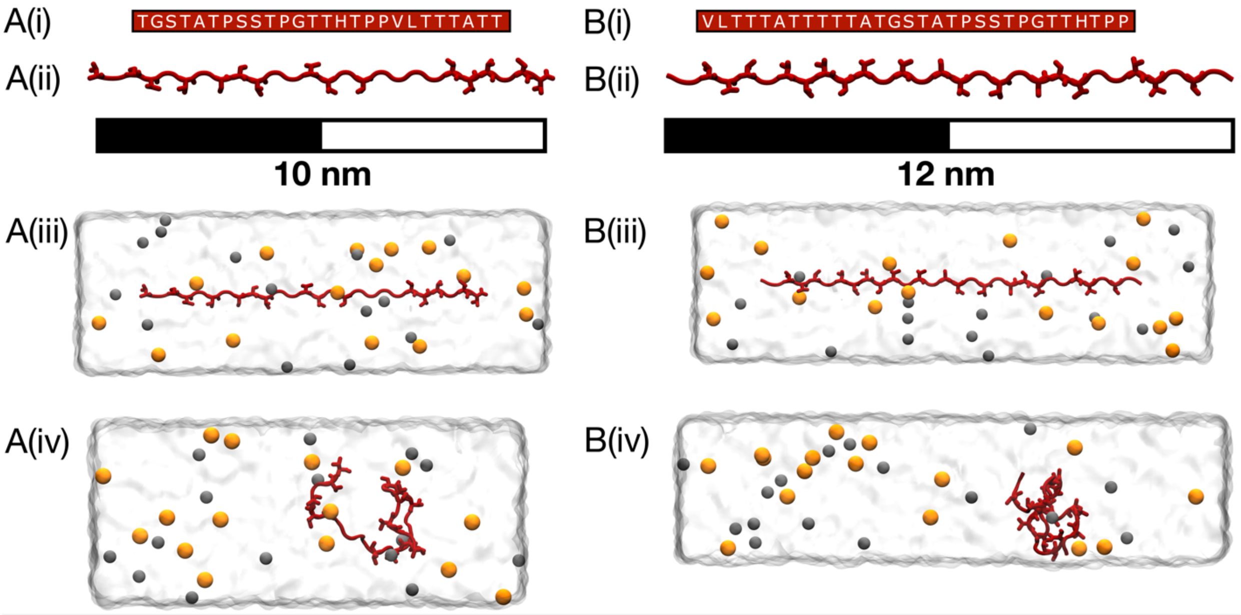

**Figure S1:** Construction of Unglycosylated mini mucin systems. (A) Mini1 (i) glycoprofile and protein sequence, (ii) pre-minimization structure, (iii) solvated and neutralized structure, and (iv) snapshot from 550ns production simulations. (B) Mini2 (i) glycoprofile and protein sequence, (ii) pre-minimization structure, (iii) solvated and neutralized structure, and (iv) snapshot from 550ns production simulations. Protein backbones are colored in dark red, glycans are colored according to SNFG color scheme, Na+ and Cl- ions are shown in orange and grey spheres, respectively.

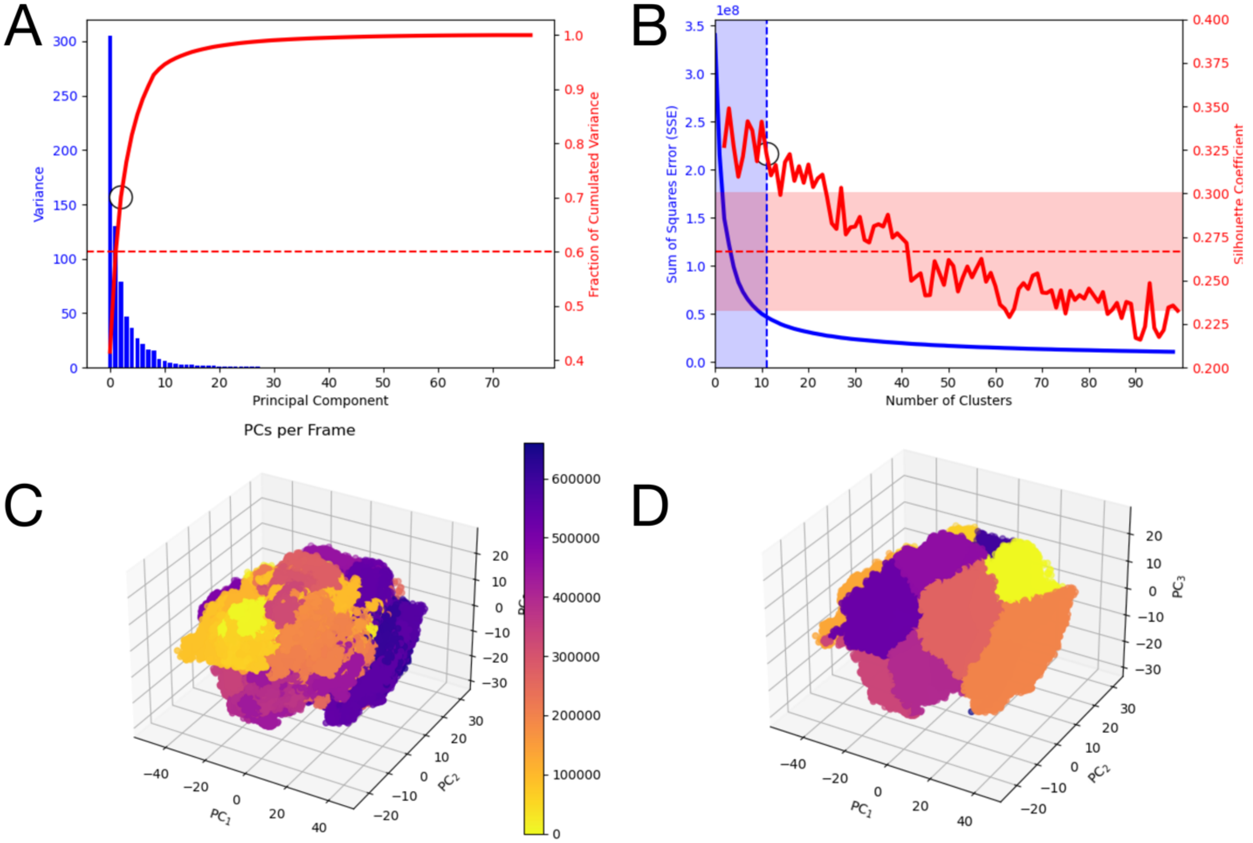


**Figure S2:** Glycosylated mini mucin clustering statistics. (A) Variance (blue bars) and fraction of cumulated variance (red line) reported for each principal component from principal component analysis of glycosylated mini mucin simulations. The top three principal components were selected as they explained > 60% of cumulated variance. (B) Sum of Squares Error (SSE, blue) and the Silhouette Coefficient (red) reported as a function of the number of clusters collected during kmeans clustering analysis. The optimal minimal number of clusters (11) was chosen which was greater than the SSE inflection point while also being greater than the average Silhouette Coefficient by one standard deviation. (C) All frames from all concatenated mucin simulations plotted on the PC1, PC2, and PC3 axes where data points are colored according to their frame number. (D) All frames from all concatenated mucin simulations plotted on the PC1, PC2, and PC3 axes where data points are colored according to their cluster numbers.

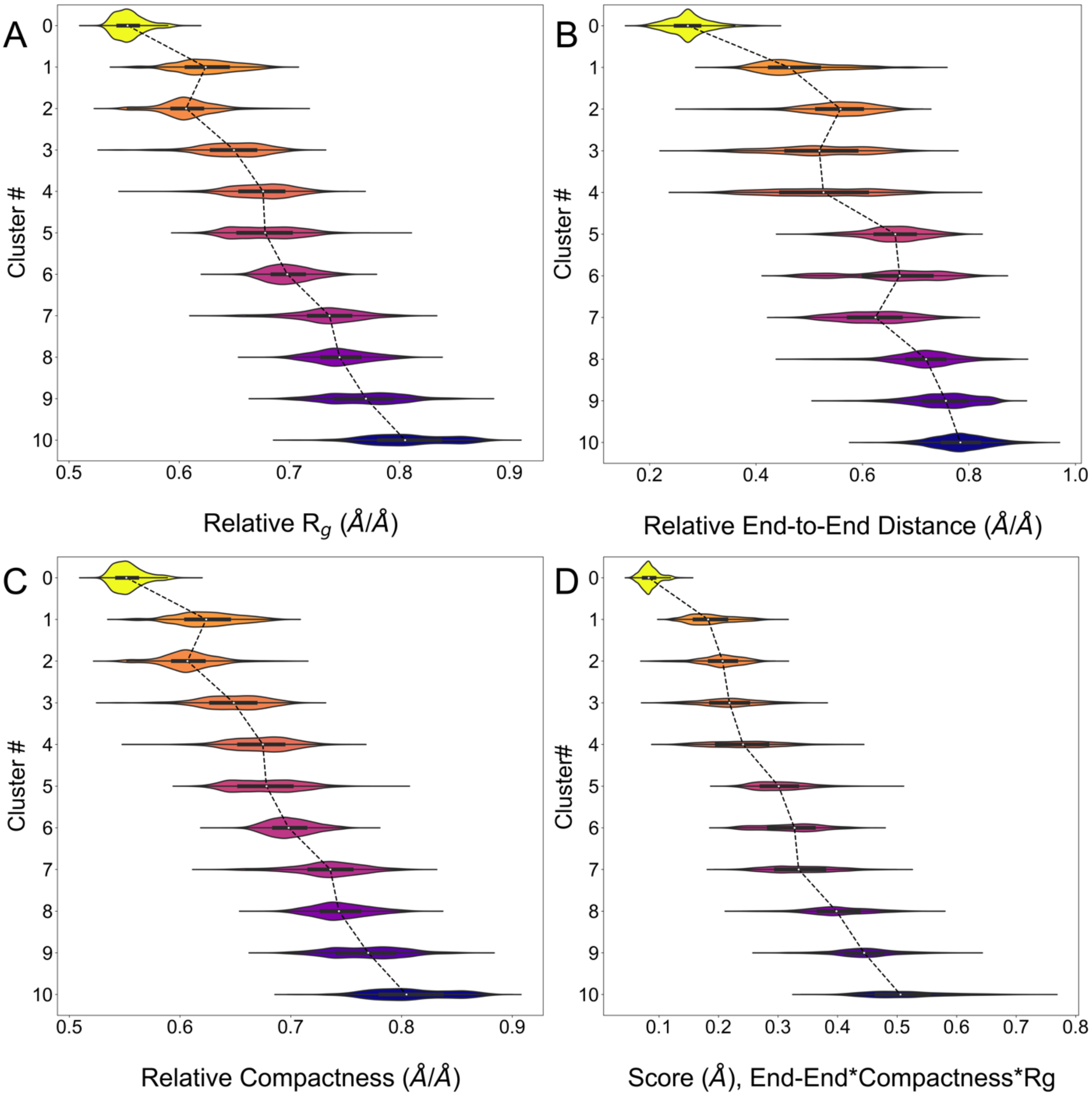


**Figure S3:** Per-cluster descriptions from the 11 clusters collected from glycosylated mini mucin simulations. Distributions are shown for (1) Relative Rg, (2) Relative End-to-End Distances, (3) and Relative Compactness. We then derived a linearity score combining all these points and have plotted (D) distributions of these linearity scores per cluster. Clusters are rank ordered according to the average (median) linearity score per cluster and all violin plots are plotted according to this median linearity score order.

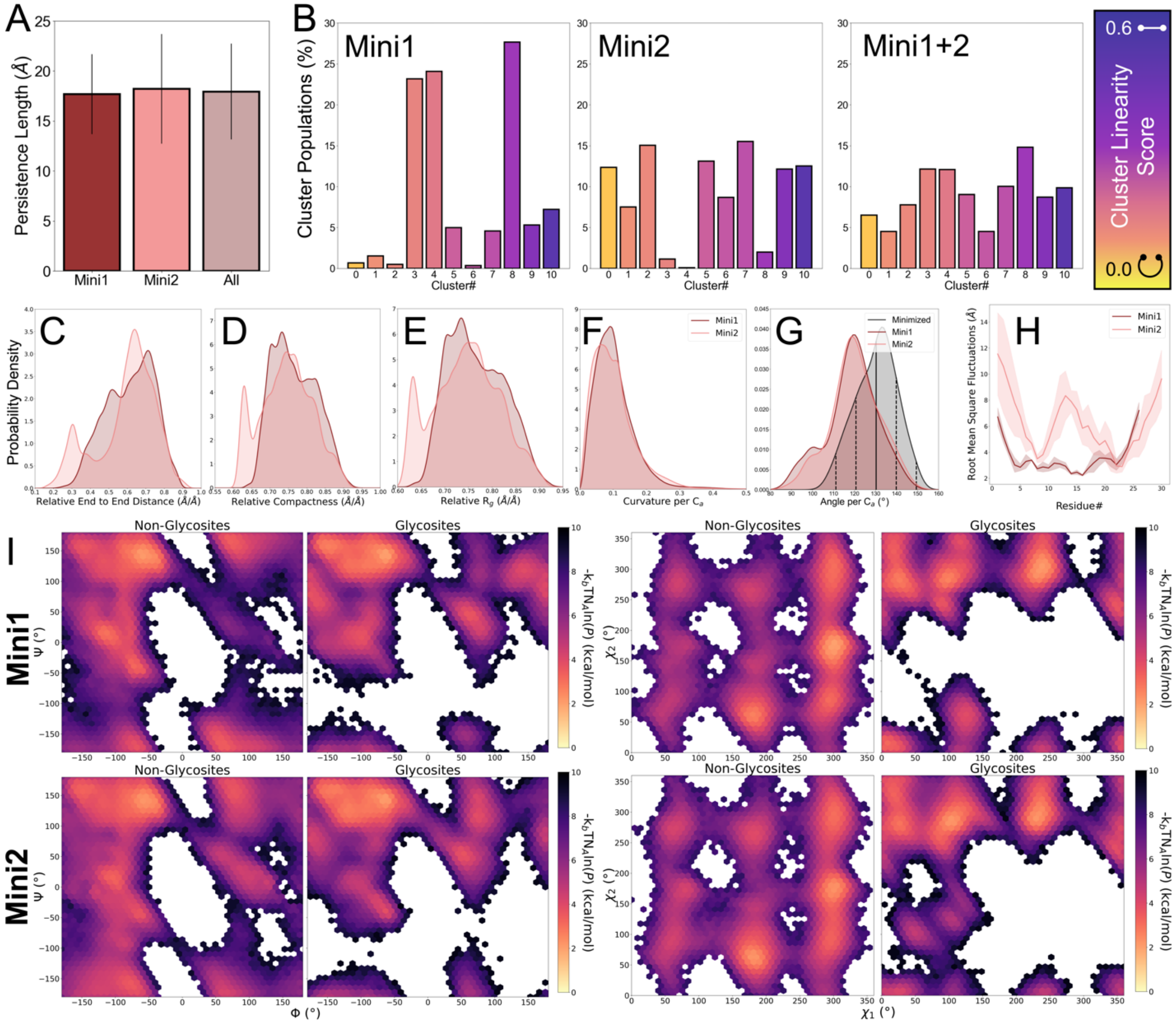


**Figure S4:** General properties from glycosylated mini1 and mini2 simulations. For glycosylated mini simulations, (A) persistence Lengths, (B) Cluster populations in percentage for Mini1, Mini2, and for all trajectories (mini1+2), probability densities for (C) relative end-to-end distances, (D) relative compactness, (E) relative radius of gyration, (F) curvature per $C_\alpha$, (G) and angle per $C_\alpha$, (H) root mean square fluctuations per amino acid, and (I) Ramachandran and Janin plots for non-glycosites and glycosites in Mini1 and Mini2. For cluster populations, bars are colored according to each cluster's linearity score. For all probability distributions shown in plots (C-G), and root mean square fluctuations (H), mini1 and mini2 distributions are shown in maroon and pink respectively. For angles per $C_\alpha$ an additional black distribution shows the distribution of $C_\alpha$ angles from minimized reference coordinate structures.

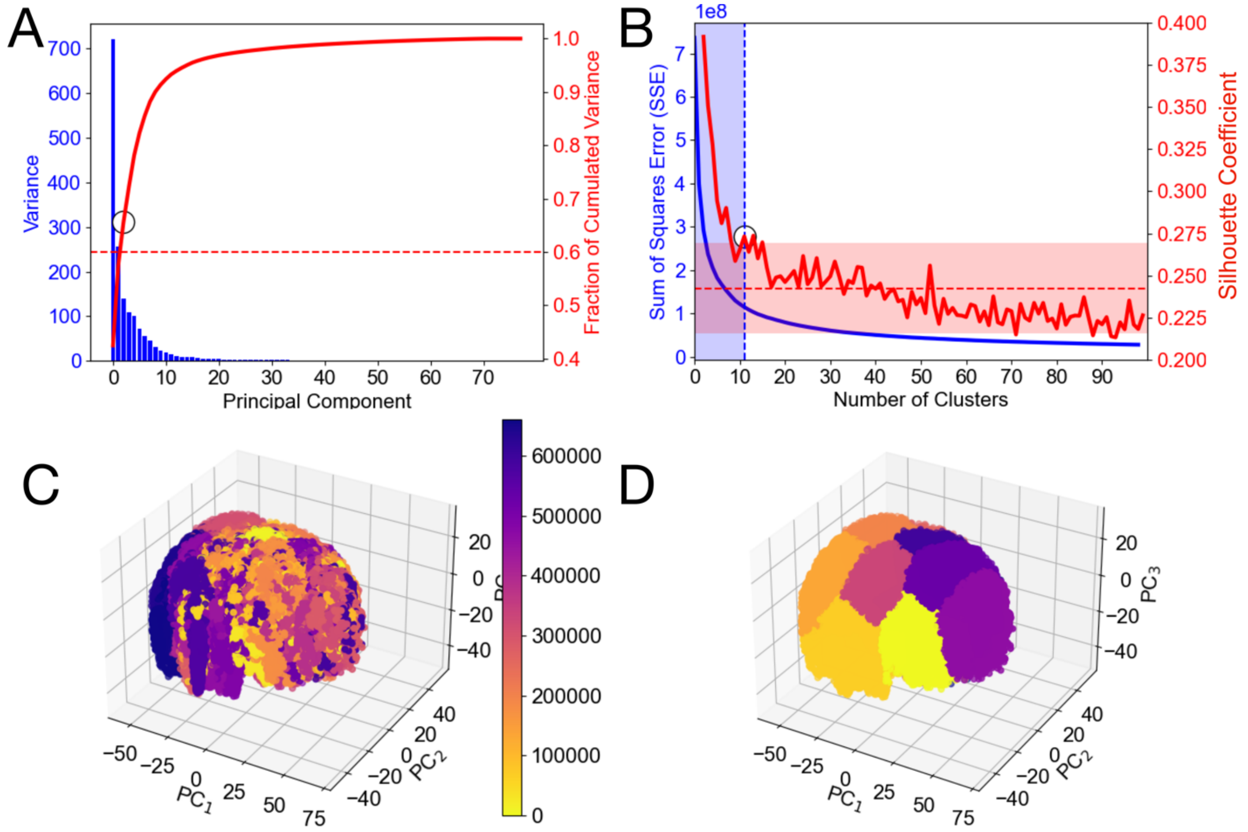


**Figure S5:** Unglycosylated mini mucin clustering statistics. (A) Variance (blue bars) and fraction of cumulated variance (red line) reported for each principal component from principal component analysis of Unglycosylated mini mucin simulations. The top three principal components were selected as they explained > 60% of cumulated variance. (B) Sum of Squares Error (SSE, blue) and the Silhouette Coefficient (red) reported as a function of the number of clusters collected during kmeans clustering analysis. The optimal minimal number of clusters (11) was chosen which was greater than the SSE inflection point while also being greater than the average Silhouette Coefficient by one standard deviation. (C) All frames from all concatenated mucin simulations plotted on the PC1, PC2, and PC3 axes where data points are colored according to their frame number. (D) All frames from all concatenated mucin simulations plotted on the PC1, PC2, and PC3 axes where data points are colored according to their cluster numbers.

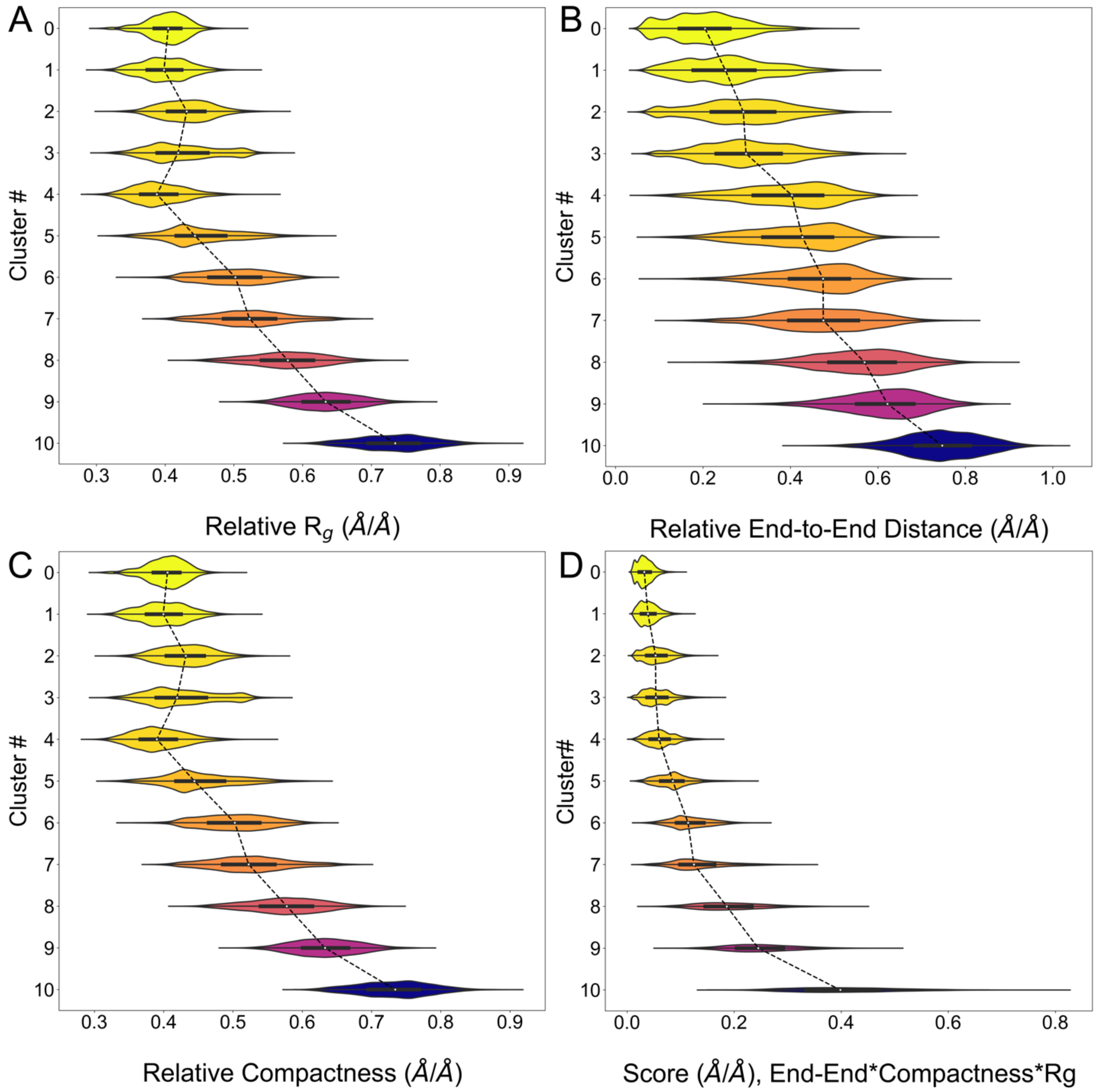


**Figure S6:** Per-cluster descriptions from the 11 clusters collected from Unglycosylated mini mucin simulations. Distributions are shown for (1) Relative Rg, (2) Relative End-to-End Distances, (3) and Relative Compactness. We then derived a linearity score combining all these points and have plotted (D) distributions of these linearity scores per cluster. Clusters are rank ordered according to the average (median) linearity score per cluster and all violin plots are plotted according to this median linearity score order.

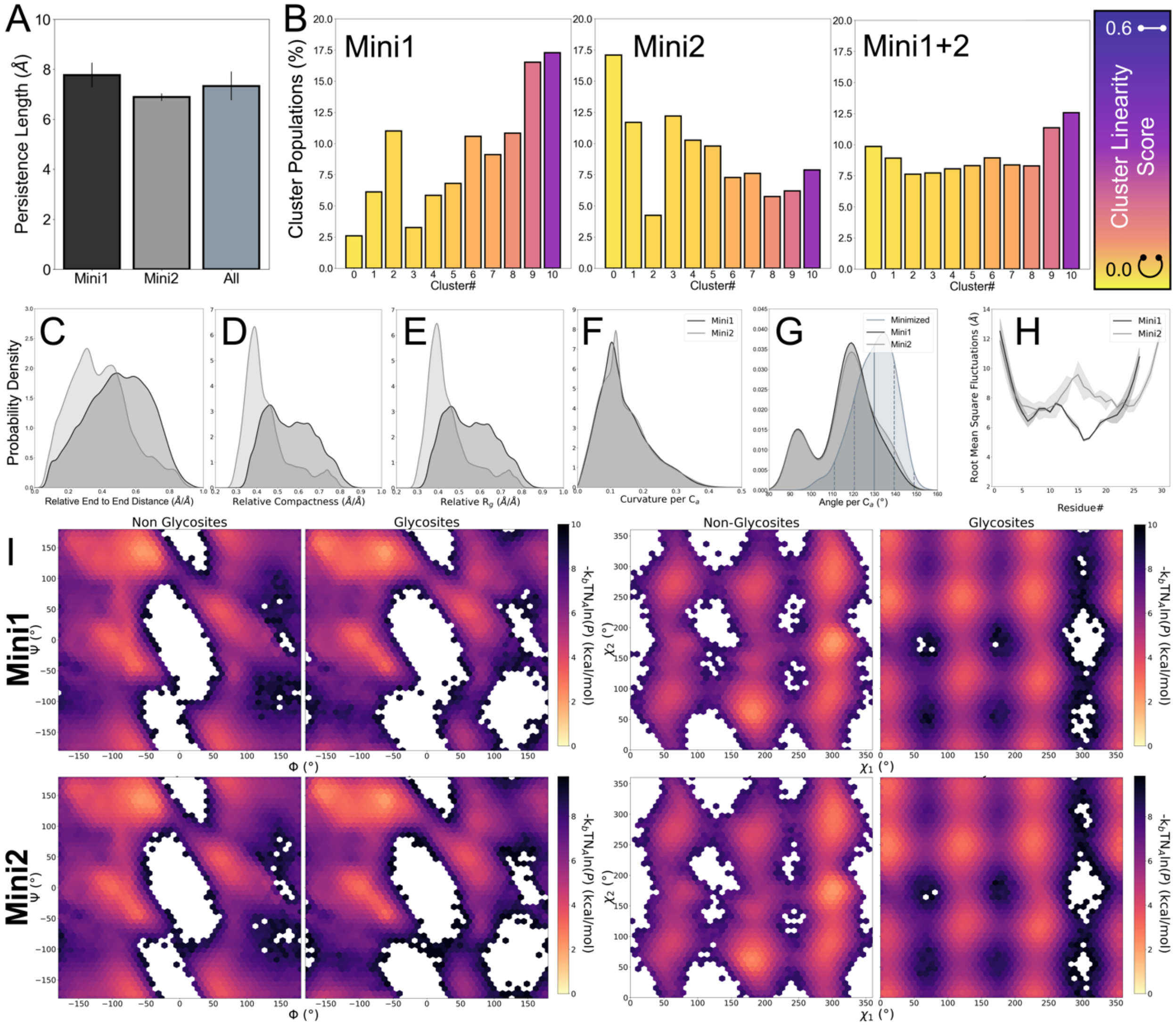

**Figure S7:** General properties from Unglycosylated mini1 and mini2 simulations. For Unglycosylated mini simulations, (A) persistence Lengths, (B) Cluster populations in percentage for Mini1, Mini2, and for all trajectories (mini1+2), probability densities for (C) relative end-to-end distances, (D) relative compactness, (E ) relative radius of gyration, (F) curvature per $C_\alpha$, (G) and angle per $C_\alpha$, (H) root mean square fluctuations per amino acid, and (I) Ramachandran and Janin plots for non-glycosites and glycosites in Mini1 and Mini2. For cluster populations, bars are colored according to each cluster's linearity score. For all probability distributions shown in plots (C-G), and root mean square fluctuations (H), mini1 and mini2 distributions are shown in black and grey respectively. For angles per $C_\alpha$ an additional slate-grey distribution shows the distribution of $C_\alpha$ angles from minimized reference coordinate structures.

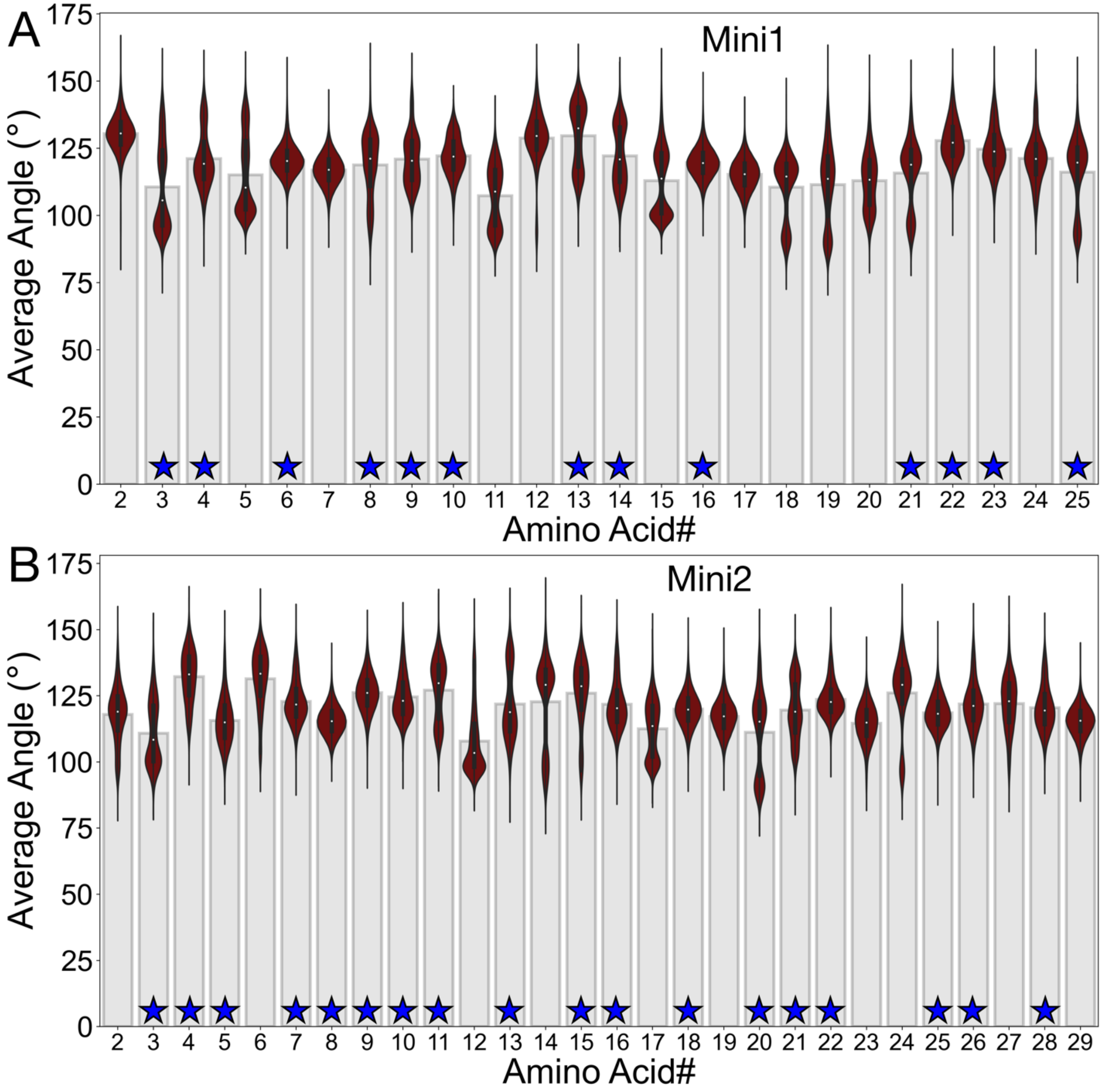


**Figure S8:** $C_\alpha$ angles calculated per residue for glycosylated (A) Mini1 and (B) Mini2 simulations. Distributions of $C_\alpha$ angles are shown in maroon violin plots over grey bar graphs plotting the average $C_\alpha$ angle per each amino acid position. Glycosylated amino acids are denoted with a blue star (position 1 in mini1 is glycosylated). Terminal $C_\alpha$ were excluded in this analysis. "Pinching" $C_\alpha$ atoms were considered to be $C_\alpha$ atoms with framewise values was less than 100°. As can be seen, there are some residues for which the distribution of $C_\alpha$ angles is bimodal with a distinct distribution lying below 100°.

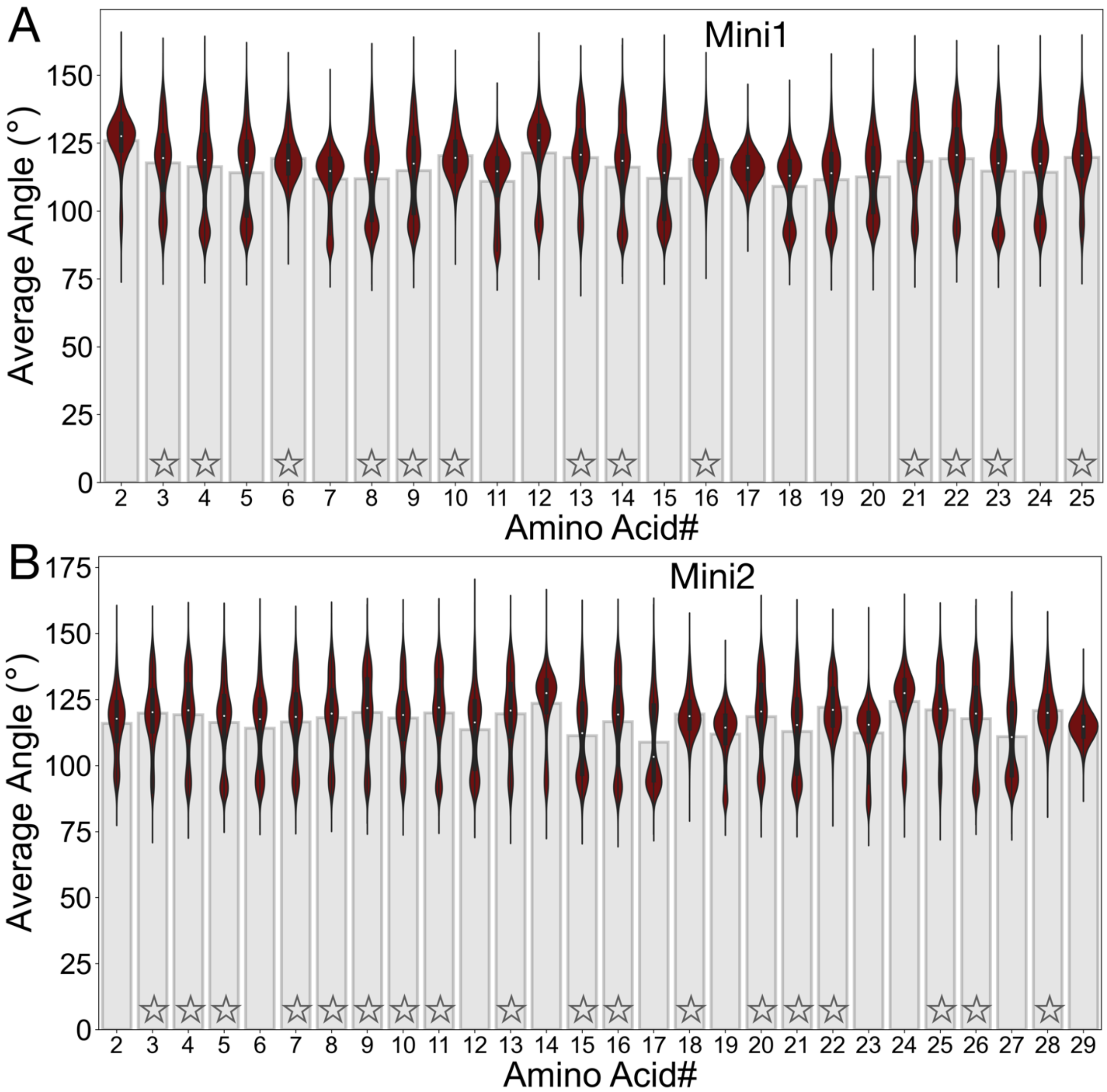


**Figure S9:** $C_\alpha$ angles calculated per residue for Unglycosylated (A) Mini1 and (B) Mini2 simulations. Distributions of $C_\alpha$ angles are shown in maroon violin plots over grey bar graphs plotting the average $C_\alpha$ angle per each amino acid position. Amino acid positions that are glycosylated in the glycosylated simulations (but are not glycosylated in these "Unglycosylated" simulations) are denoted with a grey star (position 1 in mini1 is glycosylated in glycosylation simulations). Terminal $C_\alpha$ were excluded in this analysis. "Pinching" $C_\alpha$ atoms were considered to be $C_\alpha$ atoms with framewise values was less than 100°. As can be seen, there are some residues for which the distribution of $C_\alpha$ angles is bimodal with a distinct distribution lying below 100°.

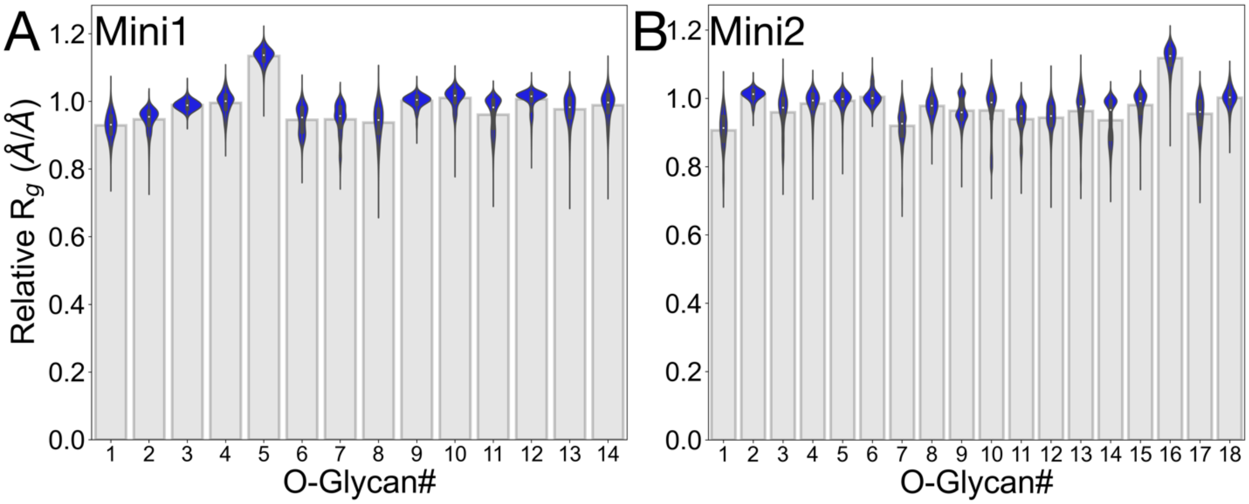


**Figure S10** Relative Rg per glycan from (A) glycosylated Mini1 and (B) glycosylated Mini2 simulations. Distributions of relative Rgs are shown in blue violin plots over grey bar graphs plotting the average relative Rg per each O-glycan.

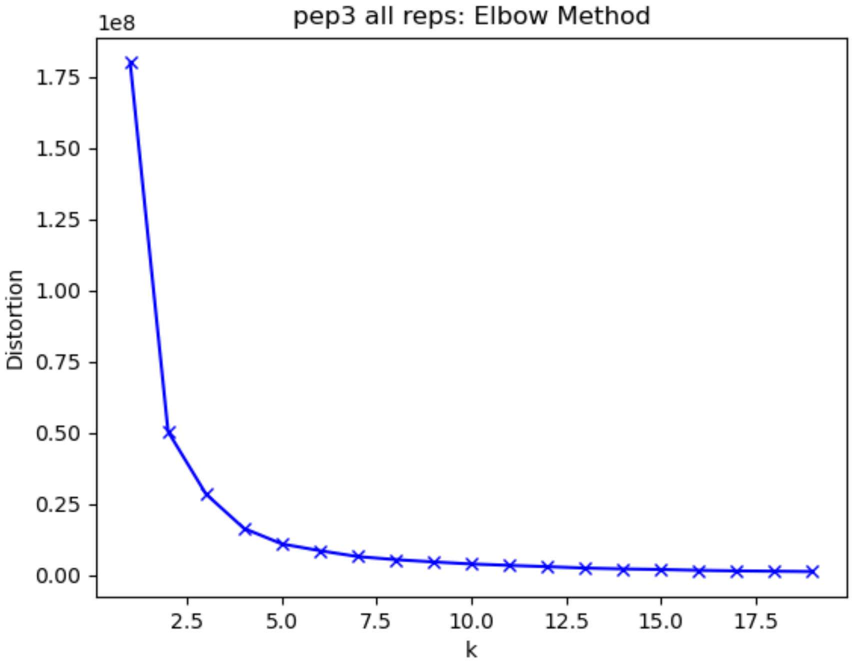


**Figure S11** Sum of squared error (SSE) for each iteration of K-means clustering over different cluster sizes. “kneed”, a scikitlearn module, was used to locate the optimal number of clusters (n=5) at the SSE inflection point.

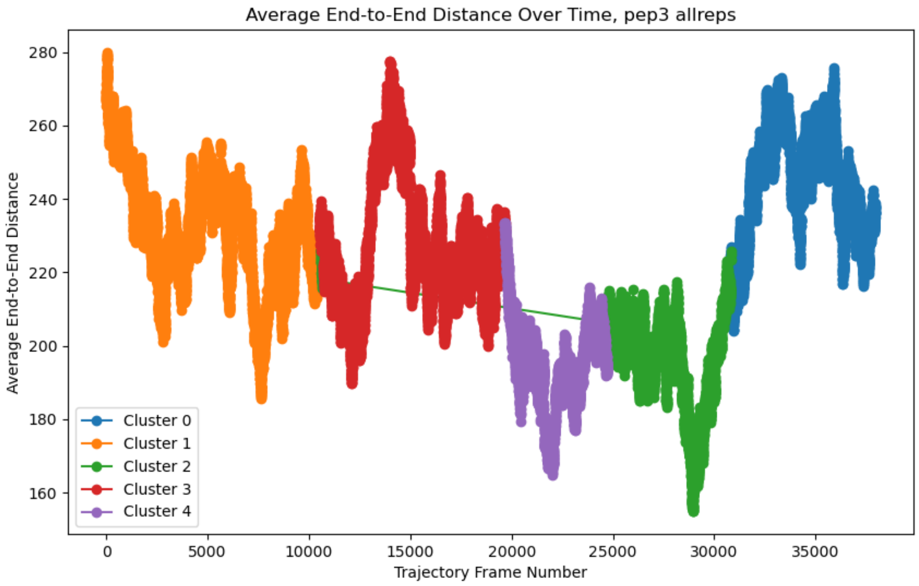


**Figure S12** Absolute end-to-end distances of glycosylated mega mucin system color coded by cluster identify. Plotting was done to ensure clusters identified with k-means clustering were of meaningful conformational differences as opposed to merely just differences in overall mucin macromolecule length.

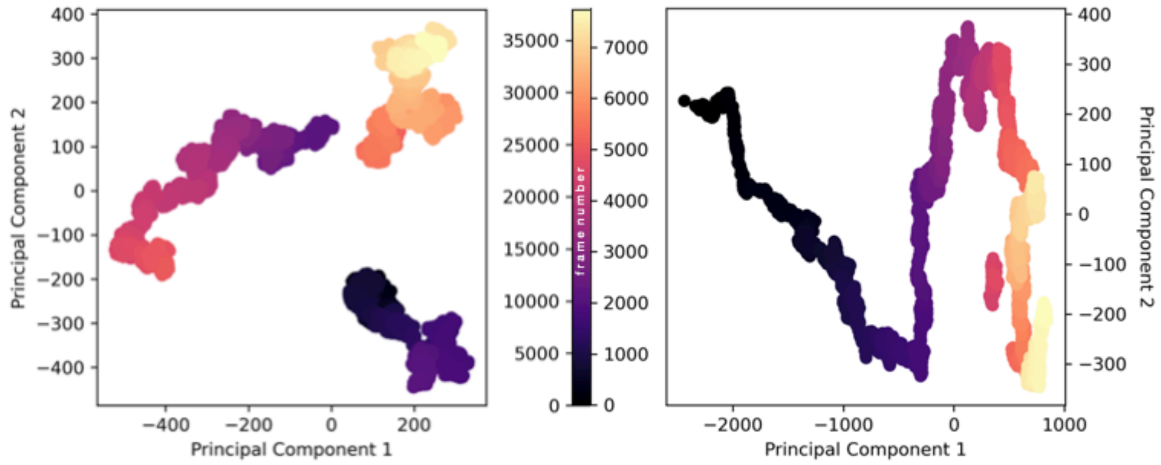


**Figure S13:** Principal component analysis of concatenated three replicas of glycosylated mega mucin system (left, to ensure description of full conformational landscape by same calculated principal components) and one replica of unglycosylated mega mucin system (right).

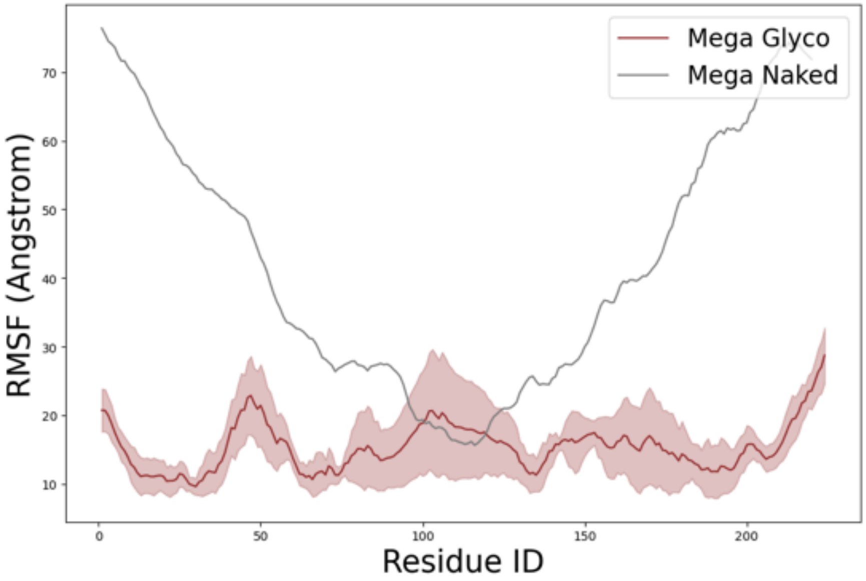


**Figure S14**: RMSFs calculated for glycosylated and unglycosylated mega mucin system.

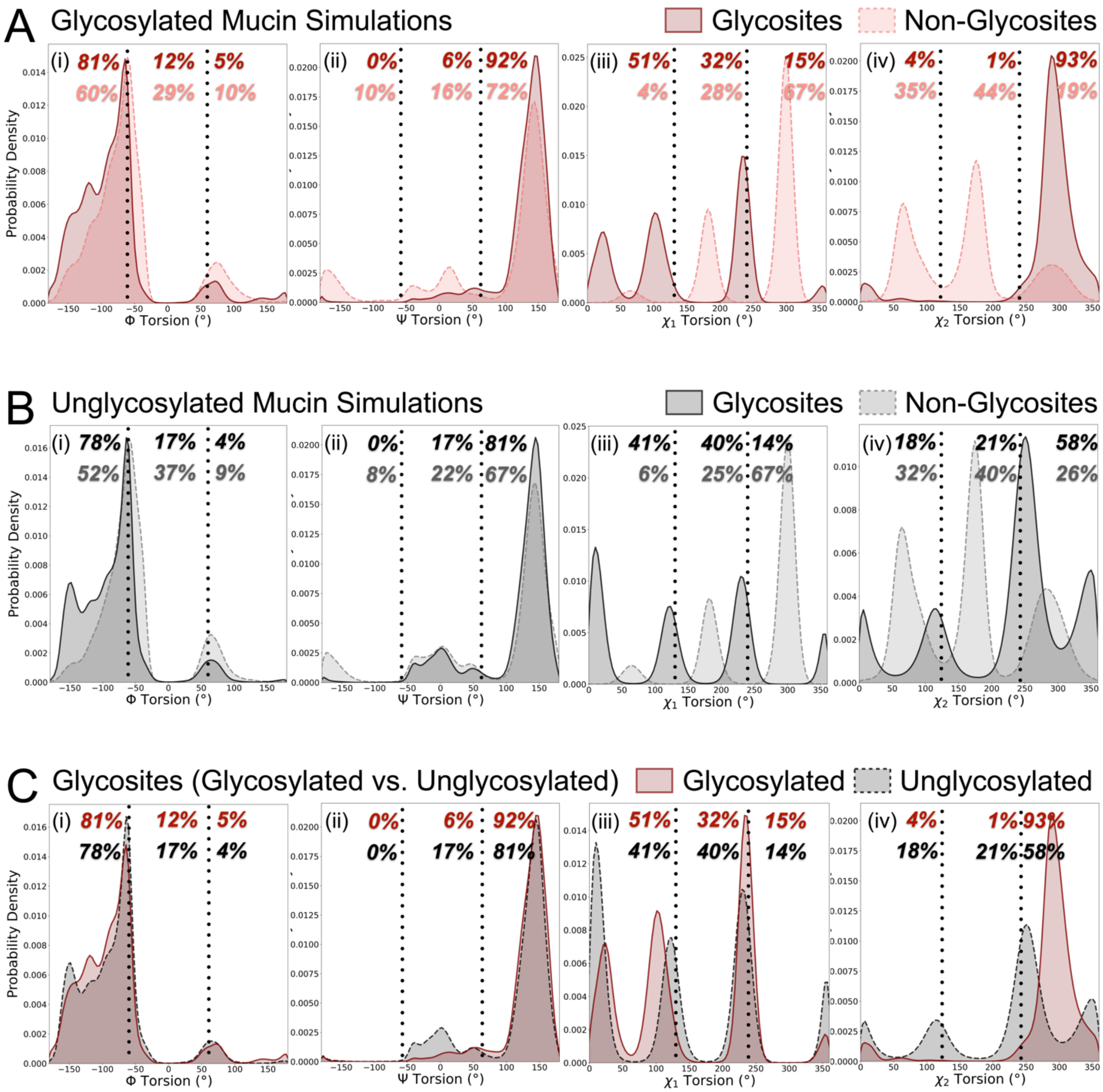


**Figure S15:** One-dimensional histograms (plotted in probability densities) for Φ/Ψ/χ1/χ2's comparing glycosites and non-glycosite residues collected from a (A) glycosylated and (B) Unglycosylated mini mucin simulations. In panel C, we compare Φ/Ψ/χ1/χ2 distributions for glycosite residues from glycosylated and Unglycosylated mini mucin simulations. Panel A: red solid and pink dashed distributions denote glycosites and non-glycosites from glycosylated mini mucin simulations, respectively. Panel B: black solid and grey dashed distributions denote glycosites and non-glycosites from Unglycosylated mini mucin simulations, respectively. Panel C: red solid and black dashed distributions denote glycosites from glycosylated and Unglycosylated mini mucin simulations, respectively.

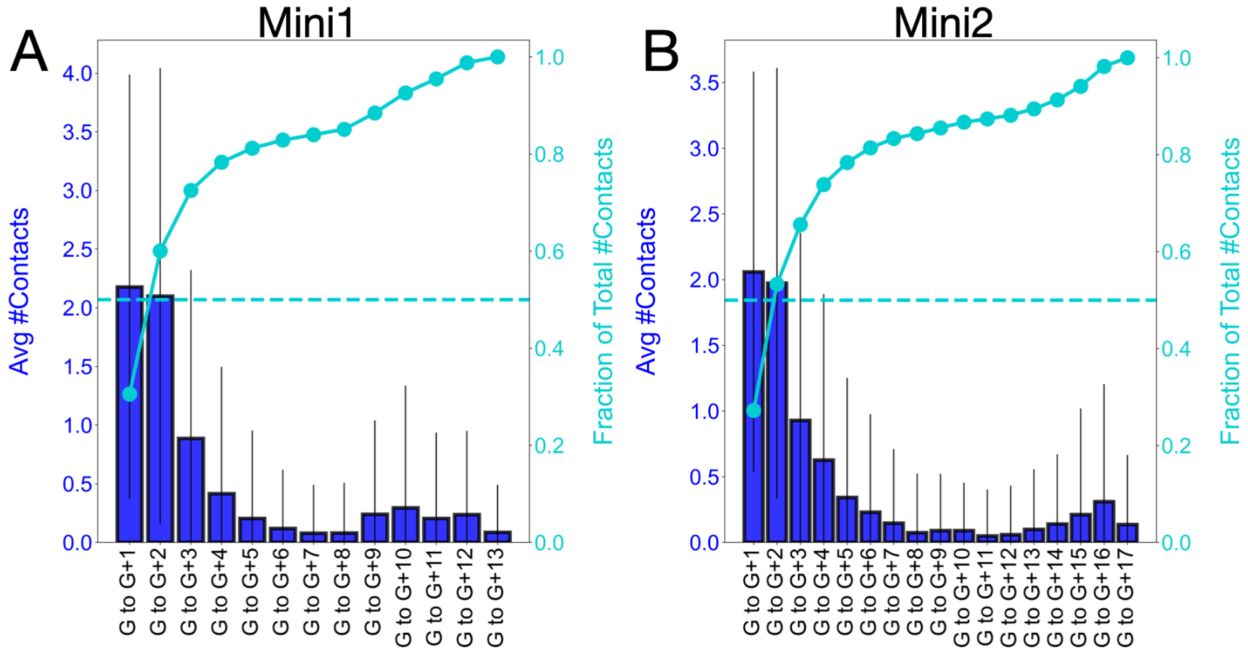


**Figure S16:** Full-size bar graphs and integration plots demonstrating the number of contacts between glycans and their neighbors for (A) Mini1 and (B) Mini2. Average number of contacts are shown in blue bars with standard deviations in those averages represented as error bars. The average fraction of total contacts is then displayed as a running total (integration) in cyan above the bar graph. It can be seen from these results that more than 50% of all glycan-glycan contacts are formed between a glycan and its two nearest neighbors (G to G+1 and G to G+2). Furthermore, 60% of all glycan-glycan contacts are formed between a glycan and its three nearest neighbors (i.e., G to G+1, G to G+2, and G to G+3).

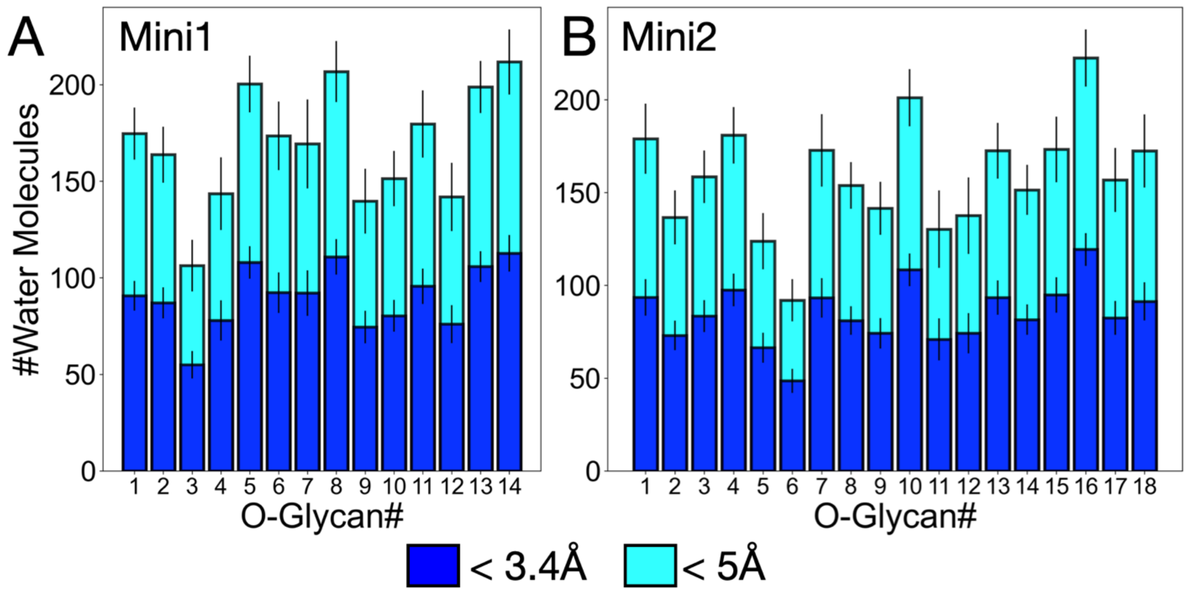


**Figure S17:** Solvation shells per glycan. First (blue) and second (cyan) solvation shell sizes per glycan, excluding protein solvation shells, for (A) Mini1 and (B) Mini2.

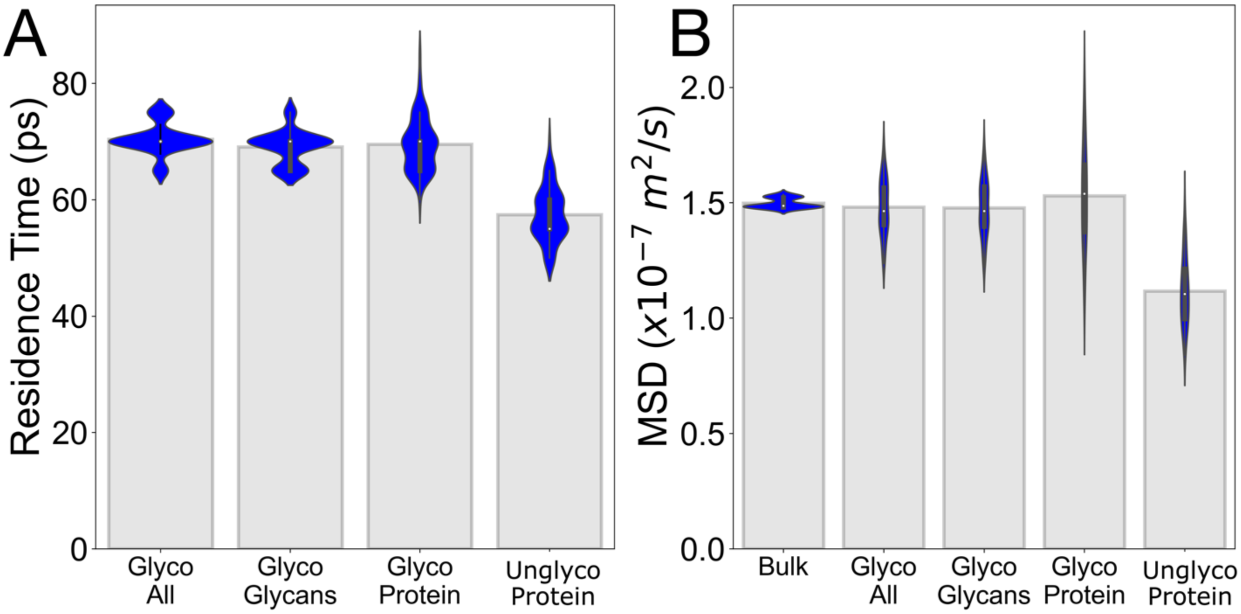


**Figure S18:** Water behavior around glycosylated and unglycosylated mini mucin models. (A) Residence time calculations for water molecules within the first solvation shell of indicated selections. Residence times were calculated as the inflection point of survival probability curves. (B) Mean square displacement values for waters within the first solvation shell of indicated selections.

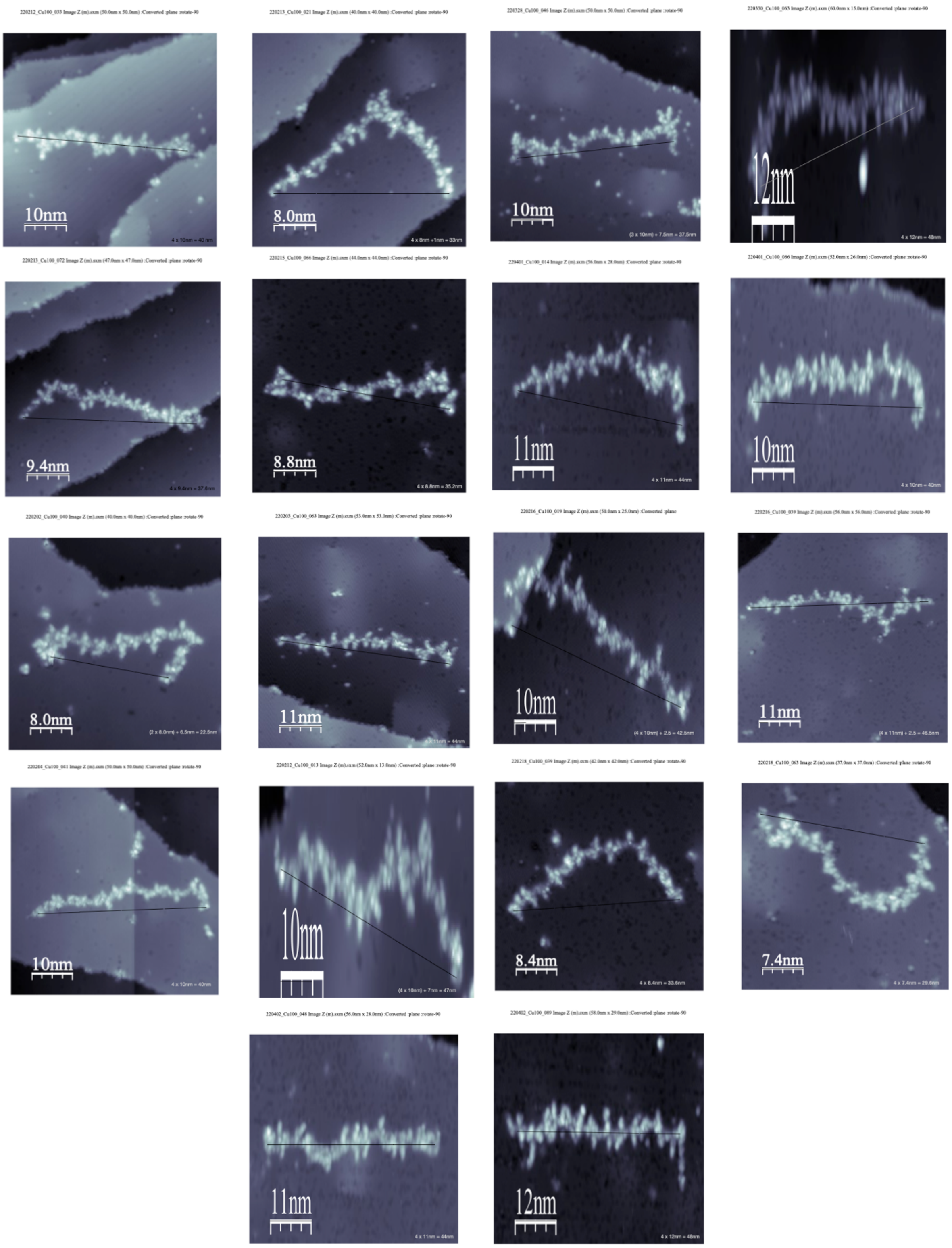


**Figure S19:** Scanning tunneling microscopy image of soft-landing electrospray ion beam deposited MUC-1 graciously provided by Anggara et al. End-to-end vectors drawn and end-to-end distances calculated at the bottom right of each image.

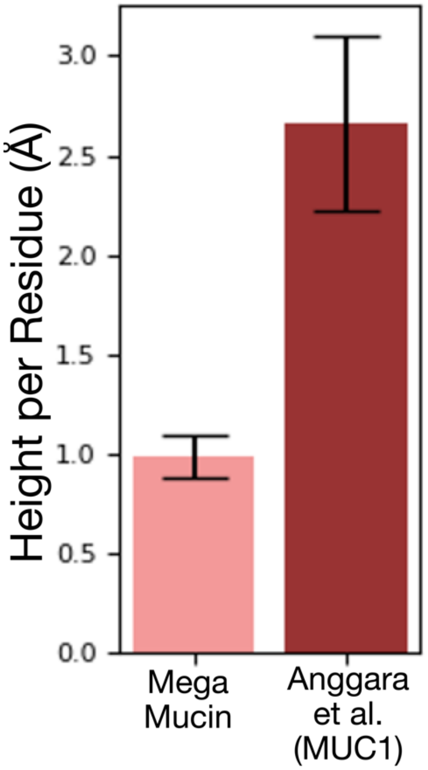


**Figure S20** Normalized measurement of height per residue in angstroms for Mega mucin and experimental STM MUC-1 images.
